# Epithelial-mesenchymal plasticity through loss of CTCF motif accessibility and protein expression

**DOI:** 10.1101/2021.06.08.447526

**Authors:** Kelsey S. Johnson, Shaimaa Hussein, Priyanka Chakraborty, Arvind Muruganantham, Sheridan Mikhail, Giovanny Gonzalez, Shuxuan Song, Mohit Kumar Jolly, Michael J. Toneff, Mary Lauren Benton, Yin C. Lin, Joseph H. Taube

## Abstract

Epithelial-mesenchymal transition (EMT) and its reversal, mesenchymal-epithelial transition (MET) drive tissue reorganization critical for early development. In carcinomas, processing through EMT, MET or partial states promotes migration, invasion, dormancy, and metastatic colonization. As a reversible process, EMT is inherently regulated at epigenetic and epigenomic levels. To understand the epigenomic nature of reversible EMT and its partial states, we characterized chromatin accessibility dynamics, transcriptomic output, protein expression, and cellular phenotypes during stepwise reversible EMT. We found that the chromatin insulating protein machinery, including CTCF, is suppressed and re-expressed, coincident with broad alterations in chromatin accessibility, during EMT/MET and is lower in triple-negative breast cancer cell lines with EMT features. Through analysis of chromatin accessibility using ATAC-seq, we identify that early phases of EMT are characterized by enrichment for AP-1 family member binding motifs but also by diminished enrichment for CTCF binding motifs. Through loss-of-function analysis we demonstrate that suppression of CTCF alters cellular plasticity, facilitating entrance into a partial EMT state. These findings are indicative of a role of CTCF and chromatin reorganization for epithelial-mesenchymal plasticity.

## Introduction

Epithelial-mesenchymal transition (EMT) is a conserved process that alters the differentiation state of a cell to drive physiological programs such as gastrulation and wound healing. During EMT, epithelial cells alter their gene expression and morphology, lose cell-cell contacts, and adopt a mesenchymal-like state (1). Because this process promotes invasion, intravasation, and resistance to anoikis, EMT is implicated in metastatic tumor cell dissemination (2–4). Recent work has contributed to a revised model of metastasis in which reversal of EMT, mesenchymal-epithelial transition (MET), is necessary for the colonization of cells which arrive at the metastatic site by means of an EMT (2,4–6). The cellular plasticity which is essential for EMT reversal is enabled by changes in genes expression and chromatin accessibility.

EMT propels cells through progressive gene expression changes and phenotypic alterations. The hallmark of EMT is the suppression of genes such as E-cadherin (*CDH1*) and epithelial cell adhesion molecule (*EPCAM*), which can be effected through networks of EMT-transcription factor proteins (EMT-TFs) such as SNAIL (*SNAI1*), SLUG (*SNAI2*), ZEB1, TWIST1, SIX1, SOX10, and FOXC2 (7–11). These transcription regulators are known to act in conjunction with epigenetic regulatory mechanisms such as post-translational modification of histone proteins (12–15) and DNA methylation (16, 17). Further key regulatory mechanisms include alternative splicing, microRNAs, and protein translation (18–24). EMT is known to be initiated by microenvironmental signals such as TGFβ, EGF, hypoxia, and tissue stiffness (25–28). Conversely, loss of such stimuli can trigger MET resulting in the re-emergence of the epithelial phenotype, re-establishment of cell-cell contacts, a decrease in migratory traits, and re-expression of epithelial-specific transcription factors such as ELF5, GRHL2, and OVOL1/2 (29–31).

Diverse partial- or hybrid-EMT states, which co-exhibit epithelial and mesenchymal traits, have been recently described and are reported to be highly plastic, and, when present in tumor cells, efficiently initiate tumor growth and predict poor patient outcomes (32–41). Distinct hybrid EMT states are likely driven by individual EMT-TFs (42, 43), which despite considerable overlap within the EMT-regulatory network, lead to distinct gene expression outcomes (11, 44). Specific transcription factor engagement can be potentiated by chromatin looping which is regulated by the CCCTC-binding factors CTCF and BORIS through their interactions with protein complexes that include cohesin and condensin (45). CTCF expression, localization, and DNA-binding activity is critical for cellular differentiation (46), yet transcription factor engagement patterns and the role of CTCF in partial EMT/MET are unclear. Therefore, we conducted phenotypic, transcriptomic, and chromatin-accessibility focused analyses of cells at progressive stages along the epithelial-mesenchymal plasticity spectrum as well as a functional assessment of the role of CTCF in epithelial-mesenchymal plasticity.

We characterized the genome-wide dynamics of chromatin accessibility at multiple timepoints during EMT and MET including the relationship between chromatin accessibility, gene expression, and cell phenotype. By staggering the time that human mammary epithelial cells (MCF10A) experienced TGFβ exposure or withdrawal, we established a series of partial EMT/MET states. Distinct phases of the repression and re-expression of the key EMT marker E-cadherin, was evident in terms of protein localization, protein expression, transcript expression, and locus accessibility. Utilizing the assay for transposase-accessible chromatin with next-generation sequencing (ATAC-seq), we report that EMT imparts a global increase in transposase-accessible chromatin while MET is marked by chromatin compaction. Transcription factor binding site (TFBS) enrichment analysis points to dynamic engagement of CTCF throughout EMT/MET. Importantly, we find that CTCF knockdown induces partial EMT. Collectively our findings indicate that activation of EMT and MET dramatically reconfigures chromatin accessibility and CTCF is a key modulator of epithelial-mesenchymal plasticity.

## Materials & Methods

### Biological resources

MCF10A cells were a gift from Dr. Sendurai Mani and were cultured as previously described (47). HMLE cell lines were also a gift from Dr. Sendurai Mani and were cultured as described previously (44). Cells were plated at 10,000 cells/cm^2^ and passaged every other day to maintain consistent densities. For TGFβ treatment, media was supplemented with 5 ng/mL recombinant human TGFβ-1 (R&D Systems; resuspended in 4 mM HCl, 0.1% BSA). MCF7, SKBR3, MDA-MB-231, and Hs-578t cell lines (ATCC) and HEK-293T cells, gift from Dr. Jason Herschkowitz, were cultured in DMEM media (Corning) supplemented with 10% FBS (Cytiva), and 1% penicillin/streptomycin (Lonza). Cells were regularly tested for mycoplasma contamination using PlasmoTest^TM^ Mycoplasma Detection Kit (InvivoGen).

### Viral transduction

FUGW-d2GFP-ZEB1 plasmid was a gift from Dr. Michael Toneff. The pHAGE-CTCF plasmid was a gift from Gordon Mills & Kenneth Scott (Addgene plasmid #116728 http://n2t.net/addgene:116728; RRID:Addgene 116728) and pHAGE-GFP was a gift from Jay Shendure (Addgene plasmid #106281; http://n2t.net/addgene:106281; RRID:Addgene_106281). Stable reporter cell lines were generated via viral transfection as described(48). pTRIPZ-shCTCF (Horizon Discovery, #RHS4696-200764825), pTRIPZ-shCtrl (Horizon Discovery, #RHS4743), pHAGE-GFP, and pHAGE-CTCF were transfected alongside pCMV-Δ8.2 and pCMV-VSVG using FuGENE HD Transfection reagent (Promega), and DMEM. Viral supernatant was clarified using a 0.45 μm filter before addition to HEK-293T cells. FUGW-d2GFP-ZEB1-GFP^+^ labeled and GFP^+^ pHAGE cells were isolated using FACSMelody^TM^ Cell Sorter (BD Biosciences). pTRIPZ-expressing cells were selected with 0.5 µg/mL puromycin.

### Cell fractionation

Subcellular protein fractionation was performed on 5 x 10^6^ cells using the Subcellular Protein Fractionation Kit for Cultured Cells (ThermoScientific) following the manufacturer’s protocol. Ten µg total protein was loaded into each well for western blotting.

### Western blotting

Cells were harvested and resuspended in RIPA buffer (Alfa Aesar) supplemented with protease and phosphatase inhibitors (ThermoScientific) and incubated in ice for 60 minutes. Lysed cells were centrifuged at 15,000 rcf for 20 minutes at 4°C and the supernatant was isolated. Protein concentrations were determined using a bicinchoninic acid assay (ThermoScientific). Proteins were separated by a 12% SDS/PAGE gel, transferred onto a 0.45 µm PVDF membrane (ThermoScientific), and probed with the appropriate antibodies. Chemiluminescence signal was obtained using ECL^TM^ Prime (Cytiva) and blot images were captured using the ChemiDoc^TM^ Imaging System (Bio-Rad).

### Flow cytometry

At the conclusion of the TGFβ time course, cells were counted and resuspended in 500 µL 1% FBS (Cytiva) in PBS with anti-E-cadherin BB700 (BD Biosciences #745965, 1:100), anti-CD44 BV421 (BioLegend #562890, 1:100), and anti-CD24 (BioLegend #311104, 1:100) and incubated on ice for 90 minutes. Following incubation, cells were pelleted and washed twice with 1% FBS in PBS and subjected to flow cytometry using a FACSMelody^TM^ (BD Biosciences).

### Reagents

The following antibodies (in 5% milk in TBST) were used for western blotting: anti-E-cadherin (#14472, monoclonal mouse, 1:1000, Cell Signaling), anti-N-cadherin (#BDB610920, monoclonal mouse, 1:1000, BD Biosciences), anti-Slug (#9585, monoclonal rabbit, 1:500, BD Biosciences), anti-CTCF (#07-729, polyclonal rabbit, 1:1000, Millipore Sigma), anti-BORIS/CTCFL (PA5-97639, polyclonal rabbit, 1:1000, ThermoFisher), anti-SMC3 (#5696, monoclonal rabbit, 1:1000, Cell Signaling), anti-YY1 (#46395S, monoclonal rabbit, 1:1000, Cell Signaling), anti-ZEB1 (21544-1-AP, polyclonal rabbit, 1:2000, ProteinTech), anti-Vimentin (#103661-AP, 1:2000, ProteinTech), anti-Actin (#612656, monoclonal mouse, 1:2000, BD Biosciences), anti-H3K27me3 (#9733S monoclonal rabbit, Cell Signaling), anti-H3K4me3 (#9751S, monoclonal rabbit, Cell Signaling), anti-H3 (#4499S, monoclonal rabbit, Cell Signaling), anti-RFP (#MA5-15257, 1:2000, ThermoFisher), anti-rabbit HRP-linked IgG secondary (#7074S, 1:2000, Cell Signaling), anti-mouse HRP-linked IgG secondary (#926-80010, 1:2000, Li-Cor).

### RNA extraction and qPCR

RNA was extracted from cell cultures using TriZol Reagent (Invitrogen) and isolated following manufacturer guidelines. 500 ng of RNA was used for cDNA synthesis. TaqMan Gene Expression primers (Applied Biosystems) were used for miR-203, miR-200c, and sno-U6. MicroRNA quantitative PCR analyses were performed using TaqMan Gene Expression Master Mix (Applied Biosystems), normalizing to sno-U6. PowerUp SYBR Green Master Mix (Applied Biosystems) and a QuantStudios5 real-time PCR machine (Applied Biosystems) was used for quantitative PCR analyses with four technical replicates per biological replicate, normalizing to *ACTB*. Signal was quantified and normalized using QuantStudios5 software (version 1.5.1) and analyzed using GraphPad Prism (version 8.4.3).

### Nanostring analysis

RNA was extracted from cell cultures using TriZol Reagent (Invitrogen) and isolated following manufacturer guidelines. 100 ng RNA was used as input for probe hybridization to a custom CodeSet. Nanostring nCounter® was used to quantify genes. nSolver 4.0™ software was used to analyze gene counts.

### Mammosphere assay

At the conclusion of the time course, 5,000 cells were plated in low-attachment 96-well plates containing 100 µL MEGM media (without BPE) (Lonza, CC-3150), 20 ng/mL FGF (Sigma Aldrich), 10 ng/mL EGF (Sigma Aldrich), 4 µg/mL heparin (Sigma Aldrich), and 1% methylcellulose. 25 µL fresh mammosphere media was added every third day. Spheres were allowed for form for 14 days.

### Wound healing assay

Cells were plated to reach 100% confluency at the end of time course. pTRIPZ-shCTCF and pTRIPZ-shCtrl cells were pretreated with 3 µg/mL doxycycline two days before wounding with a p200 tip. Measurements were made using Nikon NIS Elements Imaging Software (version 4.5) and analyzed using GraphPad Prism (version 8.4.3).

### RNA-seq Library Preparation and Sequencing

RNA was extracted from cell cultures using TriZol Reagent (Thermo Fisher) and isolated following manufacturer guidelines. Libraries were prepared using TruSeq Stranded mRNA Library Prep Kit (Illumina).

### EMT score calculation

The EMT scores were calculated utilizing the 76-gene expression signature reported (49) and the metric mentioned based on that gene signature (50). For each sample, the score was calculated as a weighted sum of 76 gene expression levels, and weights were measured based on the correlation of a particular gene with *CDH1* expression. The scores were standardized for all the samples in the dataset by subtracting the mean across samples so that the global mean of the score was zero. Negative scores calculated using this method can be interpreted as mesenchymal phenotype and the positive scores as epithelial.

### ATAC-seq Library Preparation and Sequencing

ATAC-seq libraries were generated as described (51). Briefly stated, 50,000 cells were centrifuged, resuspended in 50 µL lysis buffer (10 mM Tris, 10 mM NaCl, 3 mM MgCl_2_, and 0.1% IGEPAL CA-630), and centrifuged at 500g for 10 min at 4°C. The pellet was resuspended in transposase reaction mix (25 µL 2X TD buffer, 2.5 µL Transposase (Nextera DNA sample preparation kit, Illumina), and 22.5 µL water and incubated at 37°C for 30 min. Tagmented DNA was purified using MinElute PCR Purification Kit (Qiagen) per manufacturer’s instructions. DNA libraries were PCR-amplified using Nextera DNA Sample Preparation Kit (Illumina) using the following PCR conditions: 98°C for 30s, then thermocycling for 98°C for 10s, 63°C for 30s, and 72°C for 1 min for 12 cycles followed by 72°C for 5 min. PCR products were size-selected for 200 to 800 base pair fragments using SPRI-Select Beads (Beckmann-Coulter). ATAC-seq reads were paired-end sequenced using an Illumina NextSeq500.

### Computational Resources: ATAC peak abundance and motif analysis

Due to the similarities of both ends, one paired-end read was used for analysis. Adapter sequences were removed using cutadapt (version 1.16.6) and reads were cropped to 30 bp with Trimmomatic (version 0.36.6). ATAC-seq reads were aligned to *hg19* using Bowtie2 (version 2.3.4.2). Mitochondrial, unmapped and random contigs, and ChrY reads were excluded using samtools (version 1.9) (52) to generate filtered bam files for downstream analysis.

Tag directories were generated in HOMER from filtered bam files (53). Peaks from each replicate were called separately using HOMER ‘findPeaks [Tag Directory] -style dnase -o auto.’ Following, replicate peaks were merged by calling ‘mergePeaks -d 300 [Rep 1] [Rep 2]’ to determine common peaks. Common peaks were used for downstream analyses.

For Pearson correlation analyses, combined tag directories containing both replicates were used to quantify average peak score. Common merged peaks from each replicate were merged with other time course conditions by calling ‘mergedPeak -d 300 [Peak files].’ Time course peak files were annotated to *hg19* by calling “annotatePeaks.pl [Merged Peak file] hg19 -size 300 -log -d.” Peak scores for each condition were quantile-normalized by the preprocessCore package (version 1.44.0) in R (version 3.5.1). Quantile-normalized Pearson correlation scores were generated by R Base and visualized by the pheatmap R package (version 1.0.12).

Known motif searches were performed using HOMER (53) for 50 base pair regions excluding masked genomic regions by calling ‘findMotifsGenome.pl [Common peaks] hg19 -size 50 - mask.” Motifs with p-values > 10^-12^ were discarded. Plots representing the number of significant motifs (p-value ≤ 10^-12^) and their enrichment were generated using the ggplot2 R package (version 3.2.0).

To identify the differential peaks the HOMER function getDifferentialPeaks was used. This function was used to quickly identify which peaks contain significantly more tags in the target experiment relative to the background experiment. Here, in each pairwise comparison, Vehicle sample was considered as the background sample. The differential peaks were identified based on default parameters i.e., peaks that have 4-fold more tags (sequencing-depth independent) and a cumulative Poisson p-value less than 0.0001 (sequencing-depth dependent). Next, we annotated these differential peaks to genes and obtained the distance of these peaks from the gene transcription start site (TSS). After obtaining the annotated peaks, we looked at the peak distance from TSS together with the fold change in the expression of respective genes. The fold change for a given gene was calculated by simply taking the ratio of expression values in the two samples.

For the GIGGLE Analysis, ATAC-seq BED files were normalized using BEDTools and peaks were called using MACS2. Input peak files were queried by GIGGLE (54) against ∼47,000 CistromeDB compiled genome interval files and chromatin accessibility regulators with significant feature overlap to queried files, as quantified by similarity scores, were identified (55, 56).

### Venn Diagrams and GSEA analysis

Peak similarities were identified using mergePeaks in HOMER and annotated to *hg19* to produce the Entrez IDs for the gene promoters nearest to peaks. GSEA analysis for Molecular Signatures Database (MSigDB) hits was performed for specific peak groups. Because of the abundance of MSigDB hits, our analysis was limited to either unbiased Hallmarks-only gene sets (57), or breast and mammary-specific gene sets (containing keywords “breast” or “mammary”).

### ChIP-seq data pre-processing

ChIP-seq bed data files from Fritz et al. (58) were downloaded using SRA Toolkit (version 2.10.5). Before permutation analysis, Bedtools was used to filter out any regions overlapping a list of previously described ENCODE blacklisted and assembly gap regions for ChIP-seq (59).

### Peak enrichment analysis

For the provided CTCF ChIP-seq file, we calculated base-pair overlap between the mark and the ATAC peaks. We used a permutation-based technique to determine whether the observed amount of base pair overlap was more than expected by chance. We calculated an empirical p value for the observed amount of overlap by comparing to a null model obtained by randomly shuffling length-matched regions throughout the genome and calculating the amount of base pair overlap in each permutation. Where relevant, the p-values are adjusted for multiple testing using the Bonferroni correction.

### Oligos

The primers used for PCR amplification in this study can be found in Supporting Materials.

### Data availability

The ATAC and RNA sequencing data generated in this work have been deposited in NCBI’s Gene Expression Omnibus and are accessible though GEO series accession number GSE145851. ATAC-seq files are accessible through the UCSC genome browser [https://genome.ucsc.edu/s/kelsey_johnson1/Reversible%20EMT%20ATAC%2Dseq%20peaks].

## Results

### TGFβ induces EMT phenotypes

MCF10A human epithelial cells, derived from spontaneously-immortalized fibrocystic mammary tissue, (60) are widely utilized as model for epithelial-mesenchymal plasticity (24,61–63). To characterize progressive epithelial-mesenchymal transition (EMT), we subjected MCF10A cells to TGFβ treatment and withdrawal for varying durations (64, 65). The cells’ characteristic epithelial morphology was lost after 2 days of TGFβ treatment while a mesenchymal spindle-like morphology emerged at 4 days of TGFβ treatment (Figure 1A). Notably, withdrawal following a short-term TGFβ treatment (4 days of TGFβ) elicited a rapid return to epithelial morphology while 10 days of withdrawal was necessary for cells subjected to long-term TGFβ (10 days of TGFβ) to resolve back to an epithelial morphology (Figure 1A). To understand the reversible EMT in the context of partial and full-EMT states, we continued to use short (4 day) and long (10 day) TGFβ treatment models.

**Figure 1.**
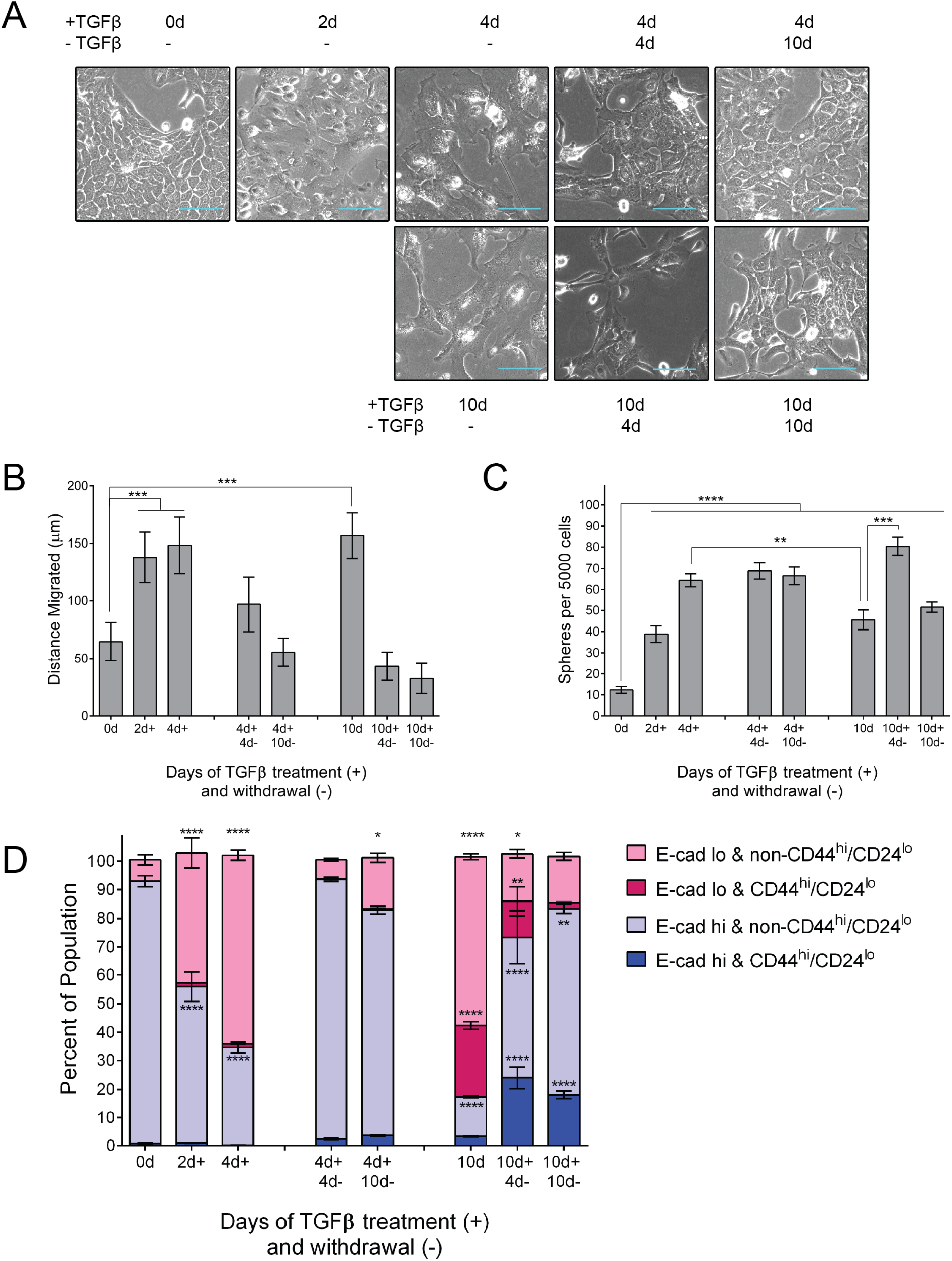
TGFβ treatment and withdrawal elicits phased cell biological changes indicative of EMT-MET. **(A)** Brightfield photomicrographs of MCF10A cells at indicated durations; TGFβ used at 5 ng/mL Scale bars = 100 µm. **(B)** Migration capacity was determined using a scratch-wound healing assay. MCF10A cells were treated as indicated prior to re-plating at confluency. The change in average gap length after 10 hours is reported. Error bars indicate s.e.m. (n = 6). Statistical significance was tested using a one-way ANOVA followed by comparison of each mean to untreated cells using a Dunnet correction for multiple hypothesis testing. **(C)** Mammosphere-formation capacity of cells treated as indicated. Following the conclusion of the time course, TGFβ-treated and -withdrawn cells were subjected to mammosphere-promoting conditions for 10 days and mammospheres ≥ 50 µm were counted. Error bars indicate s.d. (n=6). Statistical significance was tested using a one-way ANOVA followed by comparison of each mean to untreated cells using a Dunnet correction for multiple hypothesis testing. **(D)** FACS profiling of CD44, CD24, and surface E-cadherin in MCF10A cells treated as indicated. Cells were categorized as either E-cadherin-high (blue) or -low (pink) and either CD44^hi^/CD24^lo^ (lighter shading) or non-CD44^hi^/CD24^lo^ (darker shading). Error bars indicate s.d. (n=3). Statistical significance was tested using a one-way ANOVA followed by comparison of each mean to untreated cells using a Dunnet correction for multiple hypothesis testing. \**p* ≤ 0.05, \*\**p* ≤ 0.01, *** *p* < 0.001, **** *<u>p</u>* < 0.0001

We next assessed phenotypic alterations associated with EMT. As expected, cells treated with TGFβ demonstrated greater migratory capacity than control cells (Figure 1B). This effect is reversible, as TGFβ withdrawal suppressed migratory capacity (Figure 1B). Likewise, mammosphere formation capacity, an indicator of stem-like properties imparted through EMT (66), is elevated in TGFβ treated cells (Figure 1C). Intriguingly, long-term treatment imparted lower sphere-forming capacity than short-term treatment, yet cells at the initial stage of EMT reversal formed significantly more spheres (Figure 1C). These results demonstrate an enrichment in stem-like properties in partial-EMT states.

To further understand the progressive changes in stem-like phenotype, we next assayed for cell surface expression of E-cadherin, an indicator of epithelial phenotype, as well as CD44 and CD24, for which the combination of CD44^hi^/CD24^lo^ is indicative of a stem cell-rich sub-population (67). As expected, TGFβ treatment decreases the proportion of surface E-cadherin-positive cells (shown in blue) from 93.0% to 34.6% in short-term treated cells and to 17.5% in long-term treated cells (Figure 1D). Withdrawal from short-term TGFβ treatment elicited a rapid recovery of surface E-cadherin (93.7% after 4 days withdrawal) while long-term TGFβ treatment resulted in a slower recovery of surface E-cadherin expression (83.2% after 4 day withdrawal) (Figure 1D). Interestingly long-term TGFβ induces a CD44^hi^/CD24^lo^ population within the surface E-cadherin^hi^ population which comprises 3.3% of the total population, compared to 0.4% of untreated cells, (Figure 1D, dark blue). TGFβ withdrawal facilitates expansion of this population which increases to 24.0% and then 18.1%. The persistence of this population indicates a potential source of stem-like cells with partial-EMT/MET characteristics in cells undergoing MET. Overall, our TGFβ treatment and withdrawal time course reveals asymmetrical acquisition and resolution of EMT phenotypes. In particular, changes in morphology, E-cadherin localization, and migratory capacity are resolved to baseline faster than stemness-associated cell surface markers and sphere-forming ability.

### TGFβ induces gene expression dynamics

We next evaluated changes in EMT-associated gene expression by RNA-seq, qPCR, and western blotting. To measure broad changes in gene expression, we performed RNA-seq on our short- and long-term reversible EMT models and scored the gene expression based on the epithelial-specific 76-gene score (76GS) (68). Positive 76GS scores correspond to an epithelial-leaning gene expression pattern. TGFβ treatment progressively suppresses the 76GS score until TGFβ withdrawal whereupon the score progressively recovers (Figure 2A) irrespective of the initial duration of TGFβ treatment. Additionally, we assessed progression of EMT-related gene expression using a mammary cell-specific EMT signature (44). Gene expression data is plotted for 50 genes previously shown to be commonly upregulated (EMT-Up) or down-regulated (EMT-Down) by multiple EMT-TFs in HMLE mammary cells (44). In our model, EMT-Up genes increase their expression continuously throughout the time course (Figure 2B) whereas EMT-Down gene expression reaches a minimum level after 2 days of TGFβ treatment (Figure 2C). We further analyzed mRNA expression of specific genes through qPCR. *CDH1* (E-cadherin) and *CDH3* (P-cadherin), a proposed partial-EMT marker (69), show progressive downregulation which is durable through short-term TGFβ withdrawal (Figure 2D). *CDH2* (N-cadherin) and EMT-TF *SNAI2* (Slug) gene expression showed immediate responsiveness to TGFβ treatment and withdrawal (Figure 2D). On the other hand, EMT-TF *ZEB1* gene expression showed delayed upregulation (Figure 2D) while *TWIST1* (Twist) showed no significant change.

**Figure 2.**
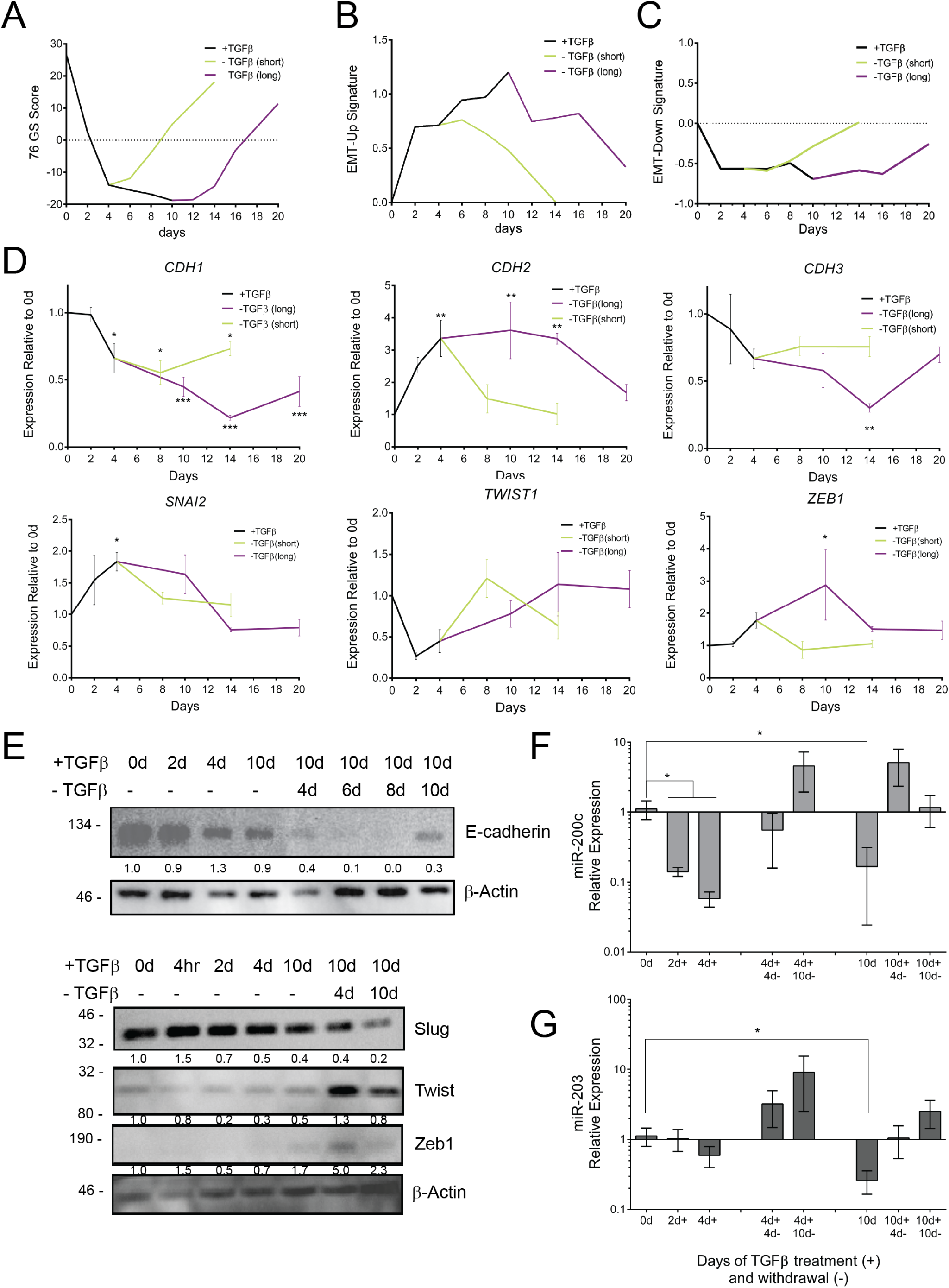
Gene expression dynamics during TGFβ treatment and withdrawal. RNA-seq data were analyzed in regard to expression of a **(A)** 76-Gene Epithelial Metric (49), **(B)** Core EMT-Up and **(C)** Core EMT-Down Gene Signatures (44). **(D)** mRNA expression of select epithelial and mesenchymal genes was determined using qPCR, normalized to untreated cells and *ACTB* and is represented as mean and s.d. (n = 3). Statistical significance was tested using a one-way ANOVA followed by comparison of each mean to untreated cells using a Dunnet correction for multiple hypothesis testing. **(E)** Western blot for epithelial (top) and EMT-TFs (bottom) in MCF10A cells treated with TGFβ for the indicated durations. Expression of **(F)** miR-200c and **(G)** miR-203 was normalized to sno-U6 and is represented as mean and s.d. Statistical significance was tested using a one-way ANOVA followed by comparison of each mean to untreated cells using a Dunnet correction for multiple hypothesis testing (n=4). *p* ≤ 0.05, \*\**p* ≤ 0.01, *** *p* < 0.001, **** *<u>p</u>* < 0.0001.

Many EMT-related genes are regulated by translation and are efficacious as proteins (24, 70); therefore, we assessed changes in protein expression for EMT markers and EMT-TFs. We observe that TGFβ induces suppression of E-cadherin and increases expression of EMT-TFs Slug, Twist, and ZEB1 (Figure 2E). Expression of the EMT-TF Slug mRNA and protein expression peaked early in TGFβ treatment while expression of Twist and ZEB1 did not reach its highest until later in the long-term TGFβ treatment model (Figure 2E). This successive enrichment of EMT-TFs may be indicative of a progressive activation of EMT-TFs wherein partial-EMT is likely driven by early-EMT TFs like Slug, as has been predicted through a modeling approach (42).

We next examined additional facets of CDH1 promoter and ZEB1 3’UTR regulatory dynamics in our model. MCF10A cells were transduced with the *CDH1* promoter linked to an RFP-encoding gene, facilitating promoter activity tracking at the single cell level (48). Commensurate with qPCR data, the *CDH1* promoter reporter remained active throughout the short-term TGFβ treatment and withdrawal timecourse. However, upon extended treatment, reporter-negative cells outnumbered reporter-positive cells, indicating promoter-induced repression (Supplemental Figure S1). Notably, the repression of the *CDH1* promoter was durable despite ten days of withdrawal. To measure post-transcriptional regulation of *ZEB1* we used a 3’UTR activity assay (48). MCF10A cells were transduced with a GFP-linked *ZEB1* 3’UTR reporter (GFP-Z1) and subjected to TGFβ treatment. GFP-Z1 expression within GFP-positive cells became elevated at two days of TGFβ treatment (Supplementary Figure S1), agreeing with the detectable ZEB1 protein and increase in *ZEB1* mRNA. Unlike CDH1, the ZEB1 3’UTR reporter readout returned to baseline upon TGFβ withdrawal (Supplementary Figure S1). Epithelial-specific microRNAs -200c (miR-200c) and -203 (miR-203) are repressed during EMT and have been reported to target *ZEB1* and *SNAI2* (71, 72). As expected, miR-200c showed a significant decrease in expression following 2 days of TGFβ treatment while suppression of miR-203 did not reach significance until completions of long-term TGFβ treatment (Figures 2F and 2G). TGFβ withdrawal elicited re-expression of both microRNAs (Figures 2F and G). Altogether these data are consistent with an early (2 days of TGFβ) suppression of miR-200c which may alleviate the repression of ZEB1 mRNA and upregulate ZEB1 protein.

### EMT and MET-induced changes in accessible chromatin regions

EMT is accompanied by reversible changes in the epigenome (47). Chromatin accessibility orchestrates dynamic gene expression by exposing or hiding genomic regulatory elements. The systematic coordination of genomic elements produces distinct epigenomic states (73). To uncover the epigenomic basis associated with TGFβ-driven reversible EMT, we performed the assay for transposase-accessible chromatin with next-generation sequencing (ATAC-seq) (51).

The extent of chromatin accessibility is highly dynamic across EMT and MET. Long-term TGFβ treatment was associated with additional ATAC-seq peaks while withdrawal from TGFβ treatment was associated with a loss of ATAC-seq peaks (Figure 3A). Dramatically, long-term treatment led to 50% more peaks than untreated cells (175,103 vs 113,680). This was reversed upon 4 days TGFβ withdrawal (10d +TGFβ, 4d -TGFβ) as the number of peaks dramatically decreased (47, 174). Though the number of called peaks fluctuate, the proportion of peaks annotated to untranslated regions (UTR), transcriptional termination sites (TTS), and exons remains stable throughout TGFβ treatment (Figure 3B). However, greater variation was evident in the proportion of intergenic and promoter-associated peaks which increase or decrease, respectively, in response to TGFβ (Figure 3B). These results suggest a broad effect of TGFβ treatment on chromatin structure.

**Figure 3.**
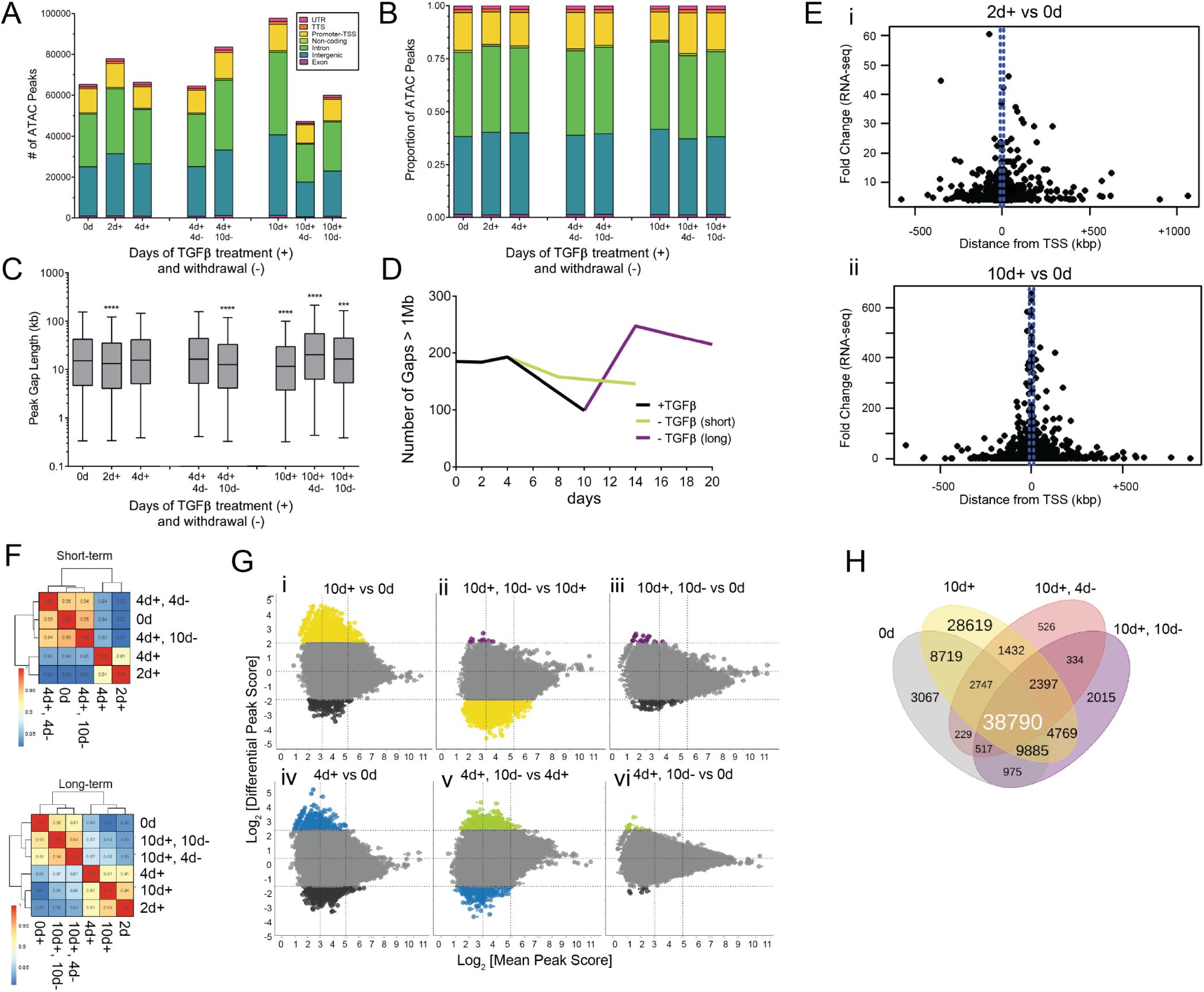
Dynamics of EMT- and MET-associated chromatin accessibility. **(A)** Number and **(B)** proportion of ATAC-seq peaks according to function of the genomic region and duration of TGFβ treatment and withdrawal. **(C)** Gap length distribution between ATAC peaks. Center line represents mean, box edges represent 25^th^ and 75^th^ percentiles, whisker ends represent 5^th^ and 95^th^ percentiles. Data were analyzed using two-way ANOVA using Tukey’s multiple comparison test vs 0d. ns = not significant, \**p* ≤ 0.05, \*\**p* ≤ 0.01, *** *p* ≤ 0.001, **** *<u>p</u>* < 0.0001. **(D)** Number of gaps between peaks that exceed 1Mb, excluding centrosome sequence. **(E)** Scatter plot showing fold-change in gene expression (RNA-seq) vs. distance from TSS for specific genes showing differential peak intensity for (i) 2d +TGFβ vs 0d and (ii) 10d +TGFβ vs 0d timepoints. Each dot represents a peak with at least 4-fold more accessibility. **(F)** Pearson’s correlation analysis on quantile-normalized log-transformed ATAC peaks. **(G)** Differential accessibility (log_2_fold change in reads per 300-bp region) among indicated conditions in long-term (i, ii, iii) and short-term (iv, v, vi) TGF-β treatment models. Colored dots represent peaks with at least 4-fold more accessibility at the indicated conditions. Black = untreated, yellow = 10d +TGFβ, purple = 10d +TGFβ, 10d -TGFβ, blue = 4d +TGFβ, and light green = 4d +TGFβ, 10d -TGFβ. **(H)** Venn diagram representing the overlap between ATAC-seq peaks common among untreated and TGFβ-treated conditions.

To further determine the genomic distribution of newly-accessible chromatin regions, we interrogated the distance, in base pairs, between annotated ATAC peaks. Generally, TGFβ treatment decreases the gap size between peaks (Figure 3C). Long-term TGFβ-treated cells had significantly shorter distances between ATAC-seq peaks (median 11,564 bp) compared to control (median 15,348 bp). Because many of the peaks unique to long-term TGFβ annotate to intergenic and intronic regions, we hypothesized that TGFβ treatment decreased the number of 1 megabase (Mb) regions without a peak—reducing the number of so-called “peak deserts”. We enumerated the number of ATAC-seq peak deserts and found that TGFβ treatment decreases the number of peak deserts from 185 in control cells to 99 in long-term TGFβ (Figure 3D). To understand how changes in chromatin accessibility associate with changes in transcription, we plotted the fold change in gene expression against distance from TSS for peaks with significantly differential peak intensity between either 2d TGFβ (Figure 3Ei) or 10d TGFβ (Figure 3Eii) and control. As expected, the strongest gene expression fold change values are associated with ATAC-seq peaks nearest to TSSs. Nevertheless, genes with significant changes in chromatin accessibility up to 500 kb on either side of the TSS also exhibit altered expression (Figure 3E). Overall, these data suggests that EMT increases chromatin accessibility across the genome with TSS distal and proximal changes affecting gene expression.

Given the diverse transcriptional dynamics observed in EMT-related genes, we next examined chromatin accessibility at select genes. We first compared peaks within epithelial-specific genes *CDH1* (E-cadherin), *CDH3* (P-cadherin), and *EPCAM* (epithelial cell adhesion molecule). Accessibility at *CDH1* and *CDH3* remained high throughout short-term TGFβ treatment as indicated by minor changes in peak profiles. These results are consistent with *CDH1* relative expression and total E-cadherin protein expression, suggesting that the lack of chromatin perturbations enabled maintenance of *CDH1* transcription and E-cadherin expression. However, in long-term TGFβ, some *CDH1* and *CDH3* peaks began to diminish at 10 days of TGFβ and continued to do so despite TGFβ withdrawal, highlighting a delayed and possibly stable chromatin alteration following EMT (Supplementary Figure 2A). These results contrast with chromatin accessibility at the EPCAM promoter, where peaks scores are more tightly associated with initial exposure to TGFβ, but which fail to return to control levels even after long-term TGFβ withdrawal (Supplementary Figure 2A). This difference in the recovery of EPCAM accessibility between short and long-term TGFβ exposure is in concordance with new findings from the Scheel lab (74) that show high EPCAM linked with the ability to enter a hybrid EMT state but low EPCAM linked with an irreversible mesenchymal state.

Accessibility patterns at mesenchymal-specific genes *CDH2*, *VIM*, *FN1*, *ZEB1*, *SNAI1*, and *SNAI2* (Supplementary Figures 2B,C) are more responsive to TGFβ than epithelial genes— increasing in accessibility and peak intensity following TGFβ treatment and resolving to untreated conditions in withdrawal. In particular, there are two regions with dense clusters of peaks near the *FN1* promoter. While peak density at the promoter proximal cluster remains elevated throughout TGFβ treatment and withdrawal, peak density at the promoter distal cluster (gray box, Supplementary Figure 2B) corresponds closely to TGFβ exposure. Similarly, *SNAI1* and *SNAI2* both show greater changes in peak height at regions distal to the coding sequence or promoter regions (Supplementary Figure 2C).

### Chromatin changes occur early in EMT and are mostly reversible

Next, to determine the relationships between timepoints in terms of chromatin accessibility patterns we calculated a correlation coefficient for each pair of samples. Because of the differences in peak distribution and score among TGFβ treated and withdrawn conditions, peaks were merged into 300 bp bins, and peak scores were quantile-normalized, prior to correlation with other time points. Expectedly, untreated and TGFβ withdrawn conditions share similarities (R^2^=0.95 and 0.93 for short-term and long-term, respectively) (Figure 3F). The greatest distinction in chromatin accessibility patterns was between untreated and long-term TGFβ treatment (R^2^=0.81) and untreated and 2 days TGFβ (R^2^=0.82). The lack of similarity in chromatin peaks between untreated and 2d TGFβ suggests that chromatin reprogramming occurs very early in EMT induction, preceding many transcriptomic changes. Interestingly, the strong correlation of chromatin accessibility patterns between 2 and 10 days TGFβ (R^2^=0.94) (Figure 3F) suggests that early chromatin accessibility alterations are sustained during EMT induction.

We next asked whether major changes in chromatin accessibility occur at regions with low, moderate, or high accessibility. Mean peak scores (MPS) were generated for pairs of samples by averaging the peak scores for those two samples. MPSs below 3 were considered “low accessibility”, while MPSs above 5 were considered “high accessibility”. We next determined the differential peak scores (DPS) for each pair of samples in order to analyze their change in accessibility. By plotting these two metrics together, we observed that major changes in accessibility (defined as a DPS above 2 or below -2) occur primarily at low and moderately accessible peaks (Figure 3G) but can occur at highly accessible peaks. In particular, a major increase in the accessibility at strong peaks is evident when comparing the long-term TGFβ treatment to untreated cells (Figure 3Gi, yellow). This is followed by a major decrease in the accessibility at strong peaks when comparing long-term TGFβ treatment to cells that have undergone a full TGFβ withdrawal (Figure 3Gii, yellow). These differential peak scores are no longer evident when comparing control to fully withdrawn cells (Figure 3Giii). Analysis of short-term TGFβ treatment indicates a similar trend to a lesser magnitude (Figure 3Giv-vi). These data indicate that long-term TGFβ treatment confers a strong increase in accessibility at highly accessible regions, which are likely to directly regulate gene expression, but also at low accessible regions, which may be more indicative of transcriptional noise.

To better understand the phased implementation of EMT-associated chromatin accessibility, we analyzed the sharing of ATAC-seq peaks between timepoints in the long-term treatment model. As expected, many chromatin accessibility regions in untreated cells are retained throughout the time course (n=38,790) (Figure 3H). However, TGFβ treatment also led to ATAC-seq peaks which either faded after TGFβ withdrawal (n=28,619) or were retained throughout TGFβ withdrawal (n=2397) (Figure 3H).

To derive the functional implications of these chromatin accessibility regions, we performed gene set enrichment analysis (GSEA) for the annotated genes closest to differential ATAC-seq peaks (74). Among the long-term TGFβ peaks unique to long-term TGFβ treatment, Molecular Signatures Database (MSigDB) revealed enrichment of gene sets involved in Hallmark EMT (p=5.5 x 10^-11^), TNF⍺ signaling (p=7.0 x 10^-13^), hypoxia (p=2.75 x 10^-6^), and TGFβ signaling (p=0.00017). This enrichment diminishes following TGFβ withdrawal. We were also interested in the overall function of the genes near persistent EMT peaks, and MSigDB hits reveal enrichment of genes involved in EMT (p=1.2 x 10^-6^), apical junction assembly (p=0.0010), and mammary stemness genes (p = 1.8 x 10^-10^) (Supplementary Figure 3).

### Dynamic transcription factor engagement during EMT and MET

We next tested the hypothesis that TGFβ modulates enrichment of transcription factor binding sites (TFBS) at accessible regions to enable partial- or full-EMT. We used HOMER motif discovery analysis to determine TFBS enrichment within our ATAC-seq peaks (53). We wished to limit our analysis to global TFBS, so we segmented the peaks to narrow 50 bp regions and identified the top 20 differentially-enriched TFBSs in comparison to untreated conditions. TGFβ increases enrichment of AP-1 (17.1% in untreated, 18.9% in short- and 18.8% in long-term) and SMAD3 (0.72% in untreated, 1.53% in short- and 1.04% in long-term) binding motifs in 50 bp segments, and their enrichment resolves to baseline levels during TGFβ withdrawal (Figure 4A). To further confirm the level of SMAD protein binding activity in our time course we measured the extent of similarity between the pattern of ATAC-seq peaks and curated patterns of genomic occupancy established for DNA-interacting factors using GIGGLE analysis (54). Limiting our comparisons to genome interval files derived from mammary or breast cancer cells, we observed that similarity scores for SMAD2/3 occupancy patterns diminished over the time course of treatment (Supplemental Figure 4A). Indeed, the SMAD2/3 occupancy patterns yielded the highest similarity scores for 2 days and 4 days of TGFβ treatment but not for 10 days of TGFβ (Supplementary Figure 4A). Patterns associated with the dimeric AP-1 transcription factor family (FOS, JUN) also yielded high similarity scores for 2 days of TGFβ (Supplementary Figure 4B-E), confirming the HOMER analysis (Figure 4A).

**Figure 4.**
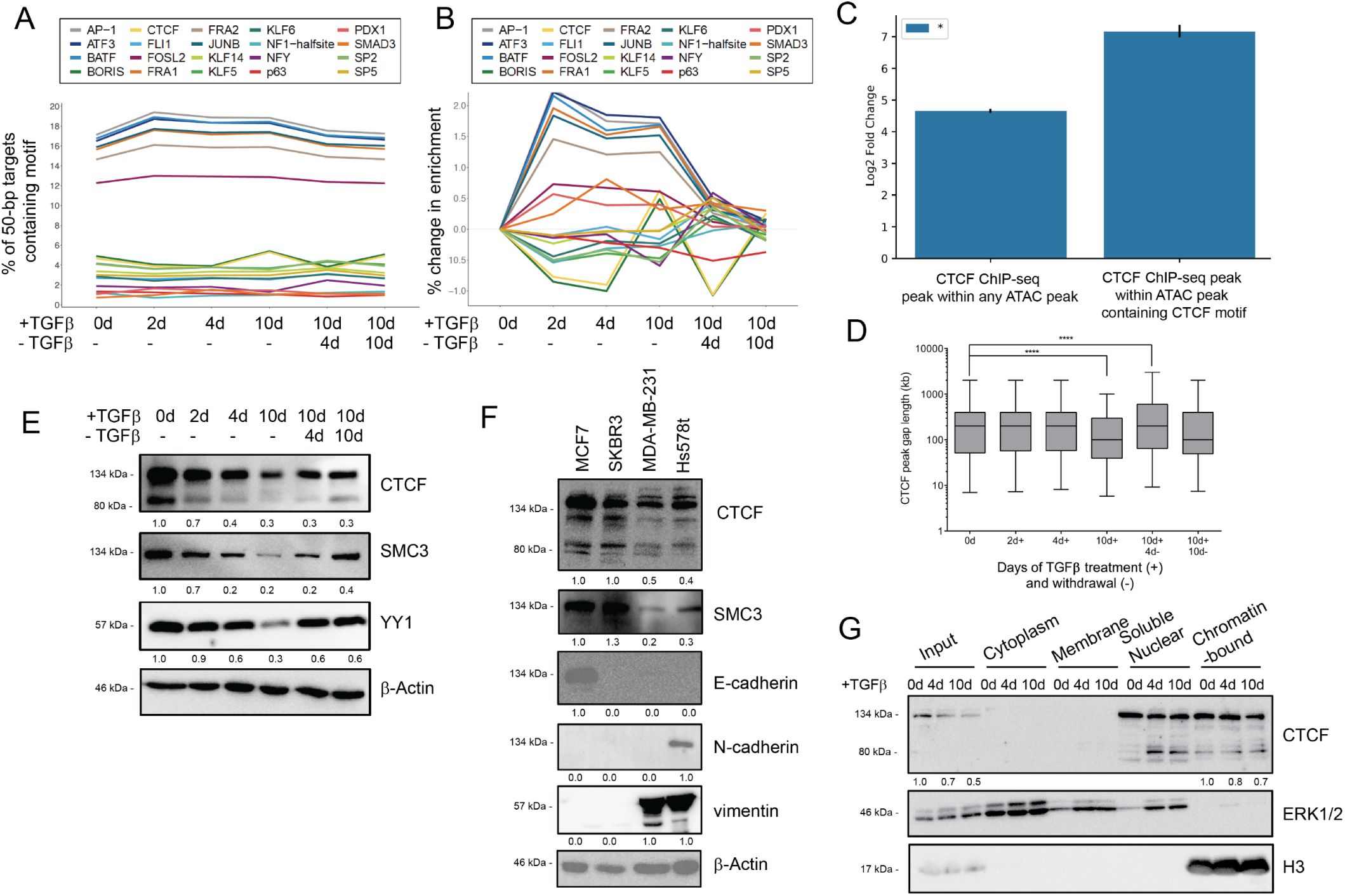
EMT-driven TFBS accessibility and CTCF expression dynamics. **(A)** Motif enrichment by percentage of 50-bp targets containing motif of the top 20 differentially-enriched motifs. **(B)** Motif enrichment percent change, compared to 0d, of top 20 differentially-enriched motifs. **(C)** The overlap between ATAC-seq peaks (in total, or with CTCF motifs) and CTCF ChIP-seq peaks from Fritz et al. (58) was compared to a random distribution model. The observed amount of base pair overlap between the all ATAC-seq peaks and ChIP-seq peaks is 2,758,444 bp, while the expected amount of overlap based on our null distribution is 110,000.50 bp (s.d. = 4894.19 bp). For the ATAC peaks with a CTCF binding motif the enrichment is greater (f.c. = 143, p = 0.001). We observe 1,728,041 bp, but expect 12,083.21 bp with a randomly distributed set of genomic regions (s.d. = 1607.12 bp) One thousand permutations were used to generate a p-value which was corrected using Bonferroni method. **(D)** Gap length distribution (in bp) between ATAC peaks containing CTCF motifs (left). Center line represents mean, box edges represent 25^th^ and 75^th^ percentiles, whisker ends represent 5^th^ and 95^th^ percentiles. Data were analyzed using two-way ANOVA using Tukey’s multiple comparison test, statistical comparisons shown in table to the right of graph, **** *<u>p</u>* < 0.0001. **(E)** Western blot for CTCF binding partners SMC3 and YY1 during long-term TGFβ treatment and withdrawal model. **(F)** Western blot for CTCF and BORIS protein expression in indicated breast cancer cell lines. **(G)** Western blot for CTCF expression in specific subcellular fractions in MCF10A cells treated with 4 or 10 days TGFβ in comparison to untreated control. H3 used as reference for quantification.

Because of the large range in TFBS enrichment amongst motifs, we also calculated the motif percent difference from untreated cells (Figure 4B). While enrichment of most motifs increased following TGFβ treatment, we observed that CTCF and BORIS motif enrichment was diminished during the EMT/MET time course, with the notable exception of long-term TGFβ cells. Indeed, at two and four days of TGFβ treatment CTCF and BORIS motif enrichment exhibited the strongest decline for any TFBS motif. TGFβ withdrawal restores these motifs to near-untreated levels.

CTCF (also known as CCCTC-binding factor) and BORIS (also known as CTCF-like or CTCFL) bind to similar cytosine-rich DNA binding motifs and exhibit some overlapping functions. Both CTCF and BORIS establish topologically-associated chromatin domains, regulate genetic imprinting, and modulate gene expression neighborhoods (75, 76). Given the unusual motif enrichment pattern, we next confirmed whether ATAC peaks considered to be enriched for the CTCF/BORIS motif overlap with validated CTCF binding sites based on ChIP-seq data in MCF10A cells by Fritz et al., 2017 (58). We observe a 143-fold enrichment for validated CTCF binding sites in our peaks with the CTCF motif (p=0.001) compared to just a 25-fold enrichment for validated CTCF binding sites in the set of all ATAC peaks (Figure 4C). These data support the notion that the peaks with a predicted CTCF-binding site correspond to *bona fide* CTCF-bound sites. The average distance between ATAC peaks with CTCF binding motifs remained stable during the initial phases of TGFβ treatment but dropped significantly at 10 days of TGFβ treatment, indicating that newly formed ATAC peaks with CTCF binding motifs are not clustering near existing peaks. Upon TGFβ withdrawal, the average distance between ATAC peaks with CTCF binding motifs stabilized back to untreated levels (Figure 4D). As with the increase in general chromatin accessibility (Figure 3C), the change in the distance between accessible regions containing CTCF motifs was consistent with novel engagement of CTCF bindings sites across the genome rather than in clusters.

Given the loss of CTCF binding sites from accessible chromatin regions during short- but not long-term TGFβ treatment, we determined how CTCF or BORIS protein expression was altered over a time course of EMT/MET. Overall, TGFβ treatment suppresses expression of CTCF protein while TGFβ withdrawal restores expression (Figure 4E). Remarkably, the expression of known CTCF binding partners SMC3 (a member of the cohesin complex) (77) and YY1 (a DNA-binding protein that forms enhancer-associated complexes with CTCF) (78) is suppressed during TGFβ treatment in MCF10A cells (Figure 4E), concurrent with CTCF. BORIS, however, exhibited a less dynamic expression pattern (Supplementary Figure 5A). In order to validate these findings in other models, we observed that TGFβ-treated HMLE mammary epithelial have decreased CTCF protein expression (Supplementary Figure 5A), and mesenchymal breast cancer cell lines MDA-MB-231 and Hs578t cells have lower CTCF protein expression than epithelial breast cancer cell lines MCF7 and SKBR3 (Figure 4F).

Considering the TGFβ-driven loss of CTCF motif enrichment in ATAC-seq peaks, we next determined if TGFβ-driven CTCF protein loss affected the nuclear and chromatin localization of CTCF. We isolated specific subcellular fractions at short- and long-term TGFβ conditions and compared the enrichment of CTCF in specific fractions. Diminished CTCF total protein was most evident in the input fraction but also observed in soluble nuclear and chromatin-bound fractions for TGFβ-treated MCF10A cells (Figure 4G) and in EMT-positive breast cancer cell lines (Supplementary Figure 5C). Although CTCF protein expression is repressed in nuclear fractions, we observe that CTCF protein is able to stably associate with chromatin in long-term TGFβ conditions which may partially explain the increase in CTCF binding motif enrichment in ATAC-seq peaks at that timepoint (Figure 4B). Surprisingly, TGFβ treatment increases the proportion of the smaller ∼80 kDa CTCF isoform, CTCF-s in nuclear fractions (Figure 4G). CTCF-s is known to antagonize binding of full-length CTCF thus altering the organization of chromatin looping (77). This smaller isoform is also evident in breast cancer cell lines and in HMLE cells exposed to TGFβ (Supplementary Figure 5B). These data indicate that expression, localization, and motif exposure of CTCF and other insulator-associated proteins are altered depending on the EMT/MET state.

We next determined if genes nearby ATAC peaks with CTCF motifs were enriched for specific functions. Comparing long-term TGFβ treatment and withdrawal, most genes adjacent to CTCF motifs were shared amongst the three conditions (n=4117) however long-term treatment had more unique genes (n=1518) (Supplementary Figure 6A). We performed gene set enrichment analysis (GSEA) for the genes uniquely-accessible in long-term TGFβ conditions and found high enrichment in gene sets regulated by chromatin bivalency (nucleosomes marked by both K27me3 and K4me3 histone H3 modifications) and neuronal development (Supplementary Figure 6B). Given the enrichment of CTCF motifs in genes related to histone bivalency, we assessed the global levels of relevant histone modifications (79). Concomitant with CTCF repression patterns, we also observe increases in H3K4me3 and H3K27me3 in TGFβ treated conditions (Supplementary Figure 6C).

### CTCF knockdown induces epithelial traits and facilitates partial EMT

We next considered the regulatory relationship between CTCF and EMT. By comparing published data (58) with our ATAC-seq data, we note verified CTCF binding sites coincident with CTCF motifs adjacent to key EMT genes. We find that CTCF motifs align to key ATAC peaks nearby *CDH3* and *CDH1* and downstream of *EPCAM* (Supplementary Figure 2A). These sites, in particular, align to peaks that diminish following TGFβ treatment. We also note several CTCF binding sites within *CDH2*, *VIM*, upstream of *FN1*, and upstream of *SNAI2* (Supplementary Figures 2B,C). The majority of the aforementioned sites align to peaks that increase following TGFβ treatment and diminish after withdrawal.

Given the suppression of CTCF during TGFβ-driven EMT and lower expression in mesenchymal breast cancer cell lines, we next investigated the functional role of CTCF expression in EMT. To do so, we generated cell lines with a doxycycline-inducible RFP plus shRNA targeting *CTCF* and probed for expression of epithelial and mesenchymal markers. Though knockdown of CTCF via doxycycline (doxy) treatment did not change cellular morphology in comparison to vehicle treated cells or doxy-treated non-targeting control (shCtrl) (Figure 5A), we confirmed doxy-induced repression of CTCF (Figure 5B). Unexpectedly, given the loss of CTCF protein during EMT, CTCF knockdown increases E-cadherin protein expression (Figure 5B). CTCF knockdown also alters the mRNA expression of EMT marker genes as noted by up-regulation of *CDH1* and basal epithelial marker *KRT5* (Supplementary Figure 7A). Concordant with CTCF’s putative involvement in H3K27me3/H3K4me3-(bivalency) marked genes (Supplementary Figure 6B), the level of H3K27me3 decreases with CTCF knockdown (Figure 5B).

**Figure 5.**
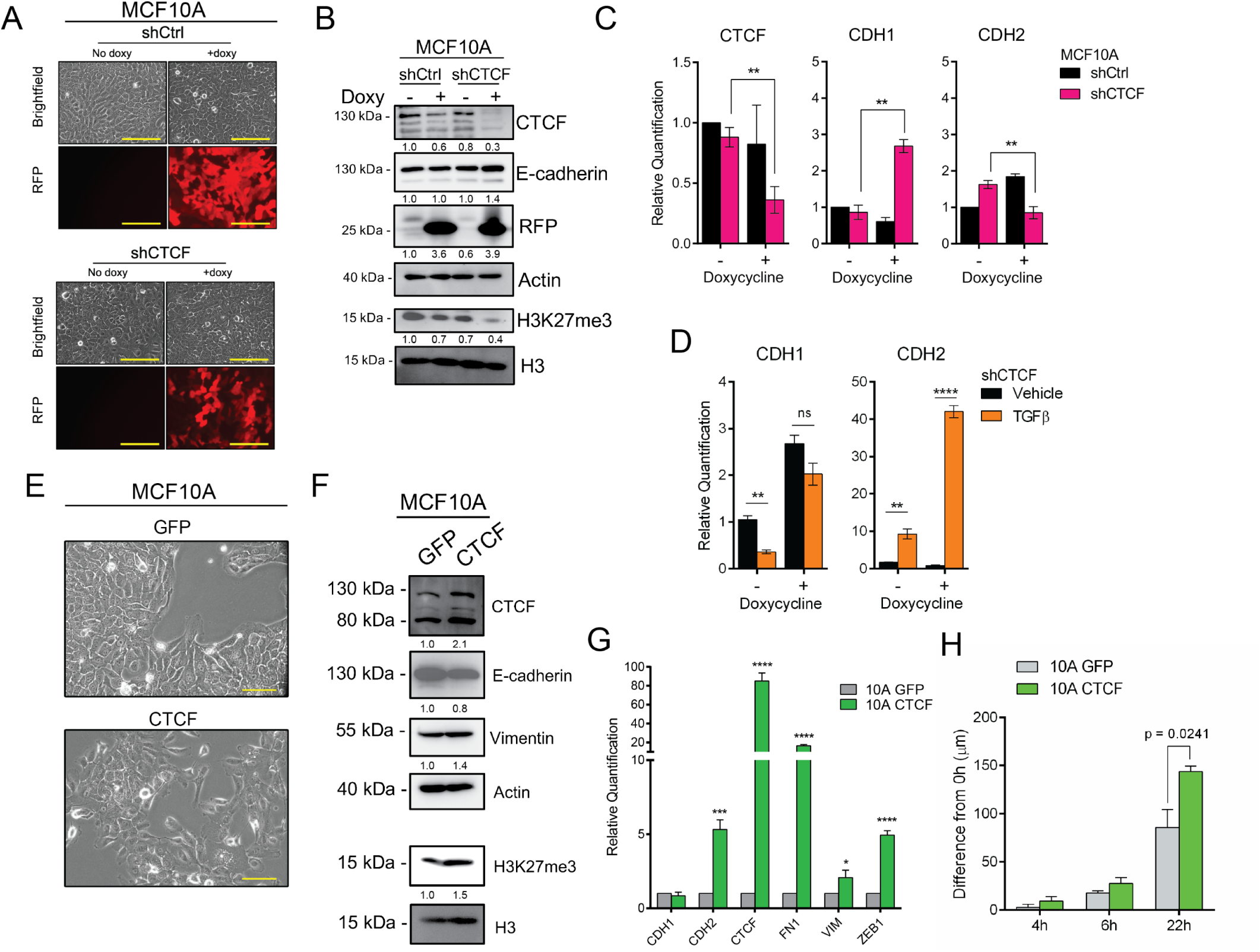
CTCF suppression and overexpression modulates epithelial and mesenchymal gene expression. **(A)** Representative images of TRIPZ-non-targeting control (shCtrl) and TRIPZ-shRNA against *CTCF* (shCTCF) MCF10A cells with and without 2 days 3.0 µg/mL doxycycline (doxy) treatment. Scale bars = 200 µm. **(B)** Western blot for indicated proteins in shCtrl- and shCTCF-expressing cells. **(C)** Gene expression measured by qPCR (normalized to *ACTB*) for indicated genes in shCtrl and shCTCF MCF10A cell lines with and without doxycycline (n = 3). **(D)** Gene expression measured by qPCR (normalized to *ACTB*) for shCTCF-expressing cells with or without 2d TGFβ treatment (n = 3). **(E)** Representative images of pHAGE-GFP (GFP) and pHAGE-CTCF-expressing MCF10A cells. Scale bars = 100 µm. **(F)** Western blots for indicated proteins in GFP and CTCF-expressing cells. **(G)** Gene expression measured by qPCR (normalized to *ACTB*) for indicated genes in GFP and CTCF-expressing cells (n = 3). **(H)** Scratch assay for GFP- and CTCF-expressing cells. The change in average gap length after 4, 6, and 22 hours is reported. Error bars indicate s.e.m., (n = 4). Statistical significance was tested using a two-tailed Student’s t-test. \**p* ≤ 0.05, \*\**p* ≤ 0.01, *** *p* < 0.001, **** *<u>p</u>* < 0.0001

Though we originally expected CTCF knockdown to induce mesenchymal traits, results of our loss-of function experiments led us to postulate that loss of CTCF slows the induction of EMT. To investigate this, we induced CTCF knockdown followed by 2 days of TGFβ treatment and assessed epithelial and mesenchymal marker expression. We limited our analysis to 2 days of EMT to capture early response of mesenchymal markers. We found that CTCF knockdown, in addition to elevating E-cadherin transcript expression, also precludes the TGFβ-mediated suppression of *CDH1* (Figure 5D). Conversely, CTCF knockdown accelerates the induction of *CDH2* transcriptional upregulation in response to TGFβ (Figure 5D). This data implicates CTCF expression in epithelial-mesenchymal plasticity of mammary epithelial cells.

### CTCF overexpression induces mesenchymal traits

We next determined the implications of CTCF upregulation on EMT. To do so, we generated *CTCF* and *GFP* over-expressing MCF10A cells. CTCF overexpressing MCF10A cells appear more mesenchymal and spindle-like (Figure 5E), suppress E-cadherin expression while upregulating vimentin protein expression, as well as the level of H2K27me3 (Figure 5F). CTCF overexpression drives transcript-level changes consistent with EMT including significant gain of *CDH2*, *FN1*, *VIM*, and *ZEB1* (Figure 5G). Moreover, CTCF-overexpressing cell lines had greater migratory capacity than controls (Figure 5H).

In conclusion, we demonstrate that TGFβ-induced EMT is accompanied by global chromatin relaxation and that TGFβ-withdrawal resolves most but not all alterations in chromatin accessibility. We show that enrichment of CTCF-binding motifs is highly dynamic across EMT/MET. Lastly, we demonstrate that CTCF expression is implicated in epithelial-mesenchymal cell plasticity.

## Discussion

EMT and its reversal, MET, are important for normal physiological processes such as development and wound healing. Together, these processes are hypothesized to endow cancer cells with metastatic abilities. Transcriptomic determination of EMT status has linked EMT to poor prognostic characterizations such as claudin-low breast cancer (44), mesenchymal glioblastoma (80), rapamycin-resistance in breast cancer (81), radio-resistance in prostate cancer (82), and immune system suppression (83). Additionally, EMT progresses into and through partial states which can assume various morphologies, gene expression, and E-cadherin profiles and exhibit greater pathogenic properties. Examination of solid tumor models has shown that cells within an intermediate-mesenchymal state are more spheroidogenic and resistant to anoikis (41). Further, cells that express both KRT14 and vimentin disproportionately contribute to metastasis (37). Despite this importance, the factors that mediate partial- and full- EMT states are not well-characterized. In this study, we have exclusively characterized the chromatin accessibly alterations and gene expression output that occur during distinct states of reversible TGFβ-induced EMT.

Herein, we show EMT and MET progress through stage-wise gene expression changes. A majority of gene expression alterations occur within 48 hours of TGFβ treatment. TGFβ treatment rapidly induces Slug (*SNAI2*) mRNA and protein overexpression, which decreases following additional TGFβ treatment into withdrawal (Figure 2D,E). These results suggest that Slug is involved in early EMT induction. *ZEB1* mRNA and protein increase after extended TGFβ treatment and the ZEB1-3’ UTR remains expressed following TGFβ withdrawal (Supplementary Figure 1). These data are similar to those recently reported by Jia et al. who find that the ZEB1 3’UTR reporter remains expressed despite prolonged TGFβ exposure and withdrawal (84). Similarly, our results corroborate the findings by Ye et al. who reported that partial-EMT states are coordinated by Slug while ZEB1 and SNAI1 promote a complete-mesenchymal phenotype (85) and data from Addison et al. who show that Slug and ZEB1 EMT-TFs both contribute to E-cadherin suppression but fail to up-regulate expression of each other (86). Our data reveal distinctions between short- and long-term TGFβ treatments, in particular, sequential activation of EMT-TFs.

We also characterized EMT/MET states in terms of surface-localized protein markers. To our surprise, we discovered that E-cadherin is lost from the membrane within 2 days TGFβ treatment despite robust total E-cadherin and *CDH1* expression. Further TGFβ treatment (4 and 10 days) yields further suppression of surface E-cadherin. Interestingly, TGFβ withdrawal following long-term treatment stimulates a return of surface E-cadherin^hi^ populations despite suppressed *CDH1* and total E-cadherin expression. Such a recovery of surface E-cadherin suggests the presence of a cytoplasmic store capable of returning to the membrane without transcriptional upregulation. This finding is in concert with past studies on E-cadherin recycling (87), including reports of the interaction between E-cadherin and late recycling endosome vesicles via Rab5 and Rab11 (70) and recent studies in MCF10A cells demonstrating that extended TGFβ treatment is necessary to induce loss of membrane-localized E-cadherin (65). TGFβ withdrawal generates a hybrid E-cadherin^hi^/CD44^hi^/CD24^lo^ population, which, given the controversial role of membrane-bound E-cadherin in metastasis (88), may have functional implications in breast cancer progression.

Through interrogation of Tn5-accessible chromatin, we identify that TGFβ treatment leads to wide-spread alterations in chromatin accessibility. Extensive chromatin alterations occur within 2 days of TGFβ treatment—revealing 22,354 more ATAC peaks and sheltering 9,678 peaks. Many newly-accessible regions have low accessibility scores which may contribute to the stochastic transcriptional noise that permits cell plasticity (89). Motif enrichment analyses reveal that TGFβ increases the enrichment for AP-1 and SMAD family binding motifs, transcription factors which have been reported to regulate EMT within accessible regions (36,37,62,90,91). Withdrawal from TGFβ induces global chromatin constriction, lower motif enrichment, resolving chromatin nearly to the untreated state. These results demonstrate that chromatin is highly-responsive to TGFβ treatment.

While most transcription factor motifs either uniformly increase or decrease following TGFβ treatment, enrichment of CTCF and BORIS binding elements declines during intermediate states. This suggests a period of chromatin re-shuffling, preparing chromatin for configurations that would promote plasticity through phenotypic states. Notably Pastushenko et al. also identified CTCF as a factor important to full phenotypic states (37). In their investigation, they isolated populations of epithelial (Epcam^+^), intermediate (CD51^-^/CD61^-^), and mesenchymal (CD106^+^/CD51^+^/CD61^+^) squamous cell carcinoma cells and subjected them to ATAC-seq (37). Motif enrichment analyses reveal that CTCF motifs were highly enriched in cells exhibiting epithelial and mesenchymal, but not intermediate, states.

In our study, we demonstrate that CTCF, a master chromatin organizer, is dynamically expressed in reversible EMT. EMT-inducing signals reduce CTCF expression while MET restores CTCF expression. We also identified that EMT-related genes (*CDH1*, *CDH2*, *SNAI2,* and *ZEB1*) contain nearby CTCF binding sites—many of which demonstrate dynamic peak profiles during the course of reversible EMT. Recently, multiple studies have implicated CTCF binding key features of cancer biology including apoptosis. For example, Kaiser et al. probed transcription factor binding sites across 11 tumor types and identified that CTCF binding sites carried high mutational loads (92). They posited that the mutations alter chromatin landscapes, replication timing, and DNA fidelity within tumors. Further, DNA methylation is well-characterized to affect CTCF binding as methylation at CTCF motifs can drive overexpression of oncogenes such as *PDGFRA* (93). Studies focusing on CTCF protein structure and DNA-binding kinetics have identified an alternatively-spliced isoform, CTCF-s, which lacks the zinc finger domains that associates with cohesin complex proteins (77). CTCF-s competes with canonical CTCF (CTCF-fl) for binding sites, consequently imposing variations in chromatin looping and gene neighborhoods (77). Indeed, we observed consistent loss of the CTCF-s isoform in cells with mesenchymal features (Figure 4E,F,G and Supplementary Figure 5B,C).

Through our investigation of the functional consequences of CTCF gain- and loss-of-function on EMT, we demonstrate that CTCF knockdown enhances the epithelial phenotype and suppresses mesenchymal markers. These findings are concordant with Zhao et al. who found that CTCF knockdown suppresses invasion and migration, proliferation, and ovarian cancer metastasis (94). Unexpectedly, we found that CTCF suppression also decreases H3K27me3, a histone modification tightly linked to EMT-induced gene bivalency (47). However, the suppression of CTCF paired with the addition of TGFβ increases *CDH2* expression without suppressing *CDH1*, suggesting that the withdrawal of CTCF from specific chromatin domains may lead to a partial EMT phenotype. Conversely, constitutive overexpression of CTCF induces a more mesenchymal phenotype including upregulation of mesenchymal markers and increased migratory speed as well as increased global levels of H3K27me3. Given the findings of Zhang et al., who determined that CTCF overexpression is linked to poor prognoses in patients with hepatocellular carcinoma (95), these findings could have considerable relevance to breast cancer. Though beyond the scope of this study, it would be noteworthy to observe how TGFβ-induced CTCF suppression affects broad 3D chromatin structure and topological domains.

In conclusion, our study demonstrates that mammary epithelial cells proceed through EMT and MET via progressive and partially reversible chromatin accessibility alterations. We reveal that EMT is marked by global chromatin loosening and MET is marked by chromatin constriction. We show that a subset of differentially-accessible ATAC peaks are enriched for CTCF binding motifs and we demonstrate that CTCF overexpression can promote an EMT-like state in mammary epithelial cells. Collectively our findings indicate that activation of EMT and MET dramatically reconfigures chromatin organization and that CTCF is a key modulator of epithelial-mesenchymal plasticity.

## Author Contributions

J.H.T. and Y.C.L. conceived, managed, and arranged funding for the project; J.H.T. and K.S.J. designed the experiments; K.S.J. performed most of the experiments and performed ATAC- and RNA-seq bioinformatics analysis; S.H., P.C., and M.K.J. provided computational analysis for RNA-seq; M.L.B. and A.M. provided computational analysis for ATAC-seq; K.S.J, G.G., and S.S. assisted with quantitative PCR data; K.S.J. and M.J.T. contributed to the Z-CAD transduced cells used in this study; K.S.J and S.M. performed western blotting; Y.C.L. provided training and guidance with bioinformatic analyses. J.H.T. and K.S.J. wrote the manuscript. All authors read and approved the final draft of the manuscript.

## Acknowledgements

We thank Dr. Dwayne Simmons for equipment sharing. We thank Dr. Michelle Nemec and the Baylor University Molecular Biosciences Center for support during the course of this work. We would like to thank Dr. Bernd Zechmann (Center for Microscopy and Imaging, Baylor University, Waco, TX) for technical support during microscopy and image analysis. This work was supported by the Collaborative Faculty Research Investment Program (30300179) and the Susan G. Komen Foundation Career Catalyst Research Grant (CCR18548469) to J.H.T. M.K.J. was supported by Ramanujan Fellowship awarded by Science and Engineering Board (SERB), Department of Science and Technology (DST), and the Government of India (SB/S2/RJN-049/2018).

---

**Supplementary Figure 1.**
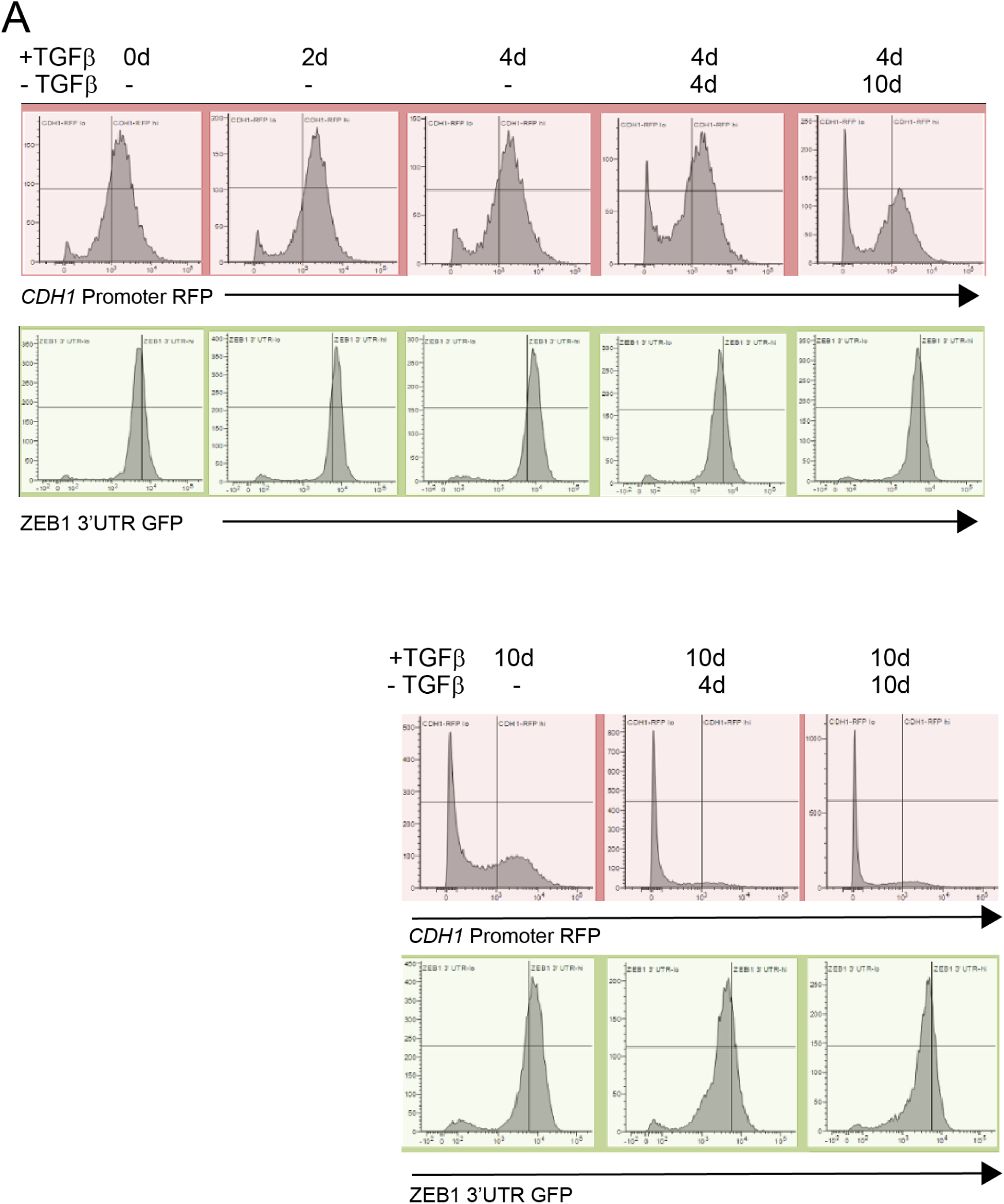
TGFβ treatment suppresses *CDH1* promoter reporter activity and increases ZEB1 3’UTR reporter activity. **(A)** FACS profiling of *ZEB1* 3’ UTR reporter and *CDH1* promoter reporter activity. FACS quantification for Z1-GFP reporter (green panel), *CDH1*-RFP reporter (pink panel).

**Supplementary Figure 2.**
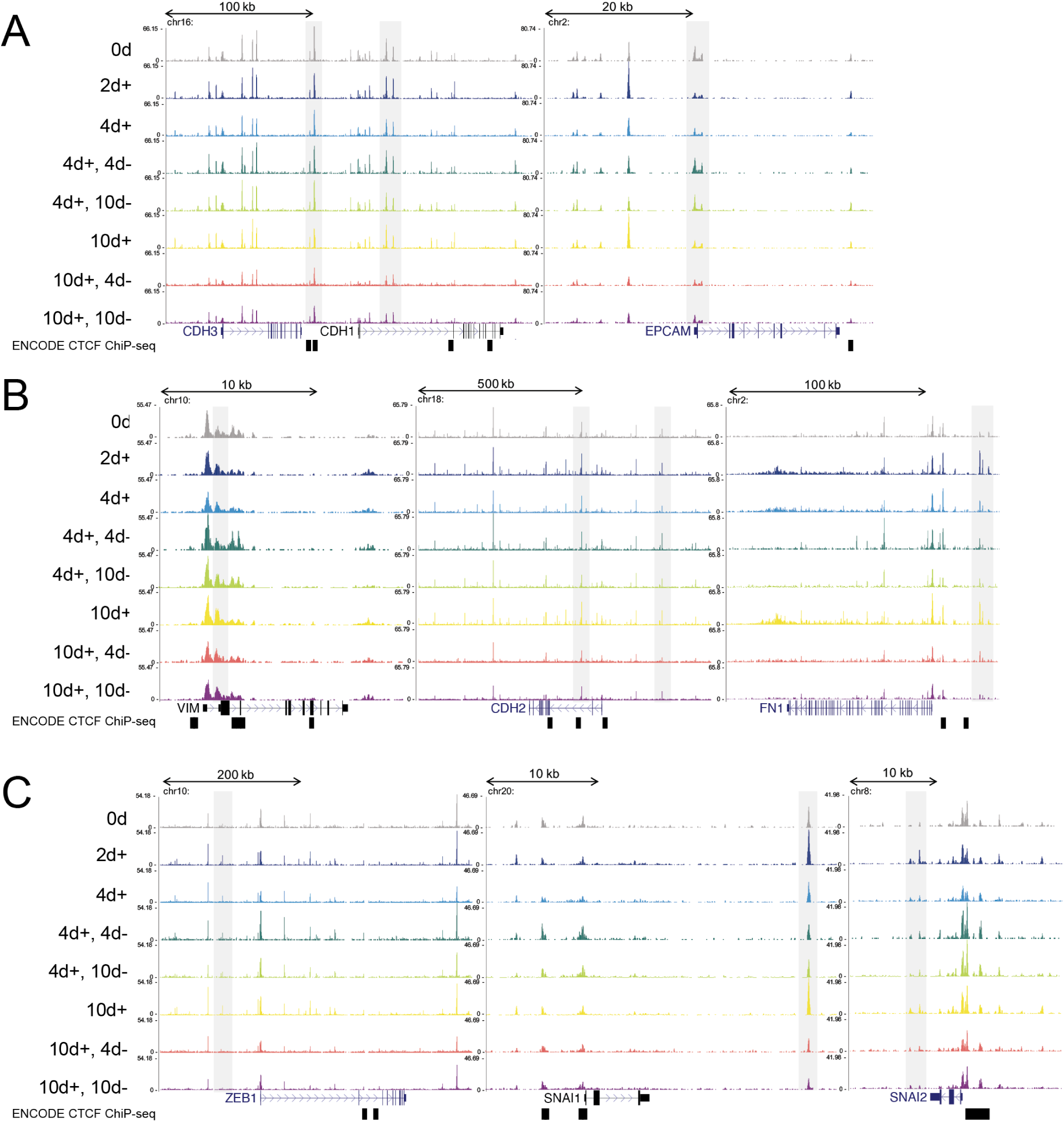
Dynamics of TGFβ-induced and TGFβ-withdrawn chromatin accessibility. ATAC-seq peak profiles of select **(A)** epithelial (*CDH3, CDH1,* and *EPCAM*), **(B)** mesenchymal (*VIM, CDH2,* and *FN1*), and **(C)** EMT-TF (*ZEB1, SNAI1,* and *SNAI2*) genes at indicated short-term (top) and long-term (bottom) TGFβ treatment models (n=2). Black rectangles below genes refer to ENCODE CTCF ChIP-seq peaks in human mammary epithelial cells (HMEC).

**Supplementary Figure 3.**
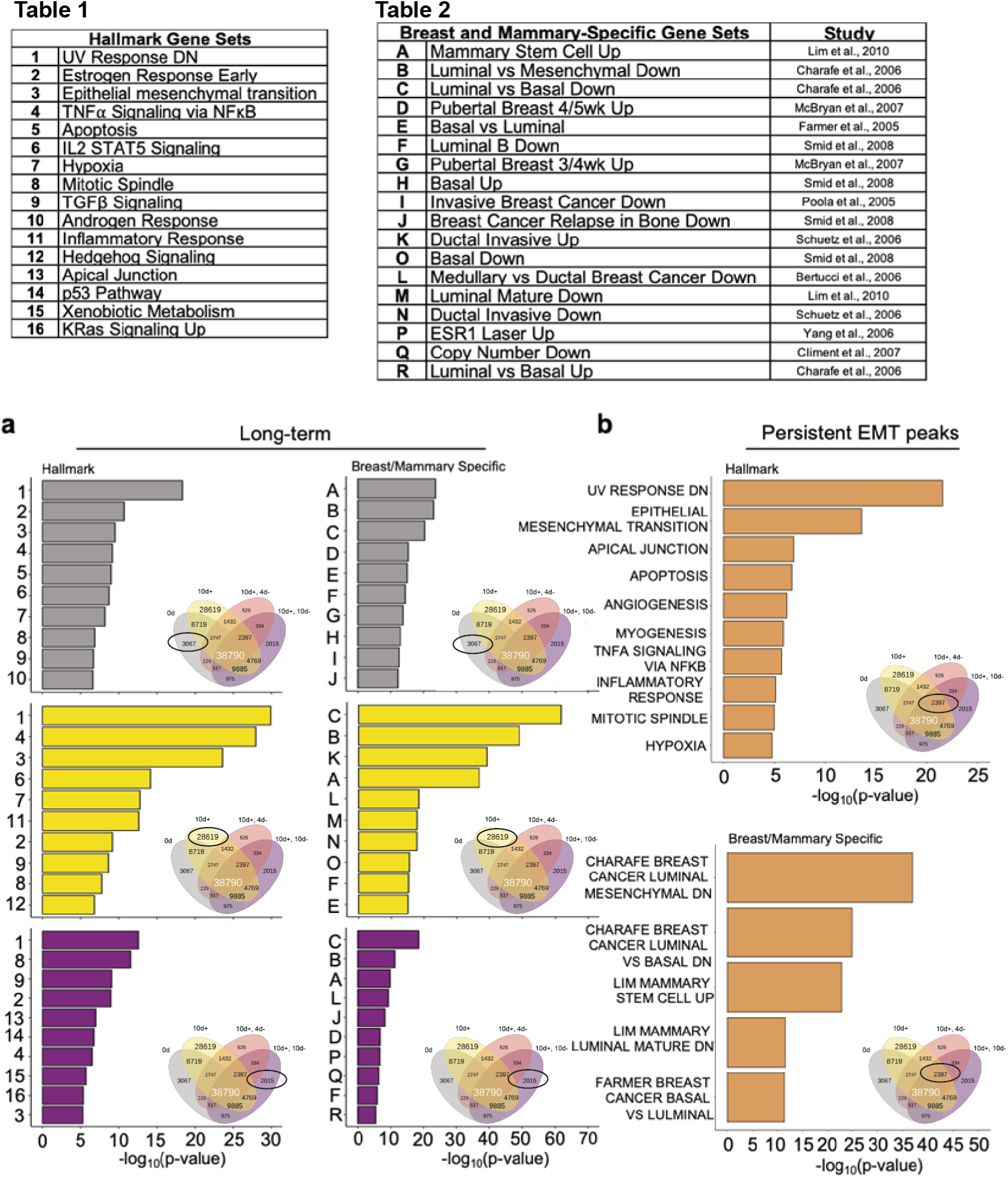
EMT, mammary basal cell, and stemness gene sets are enriched in TGFβ-induced and persistent peaks. (Table 1) Number key for GSEA MSigDB hits for Hallmark gene sets. (**Table 2)** Letter key for GSEA MSigDB hits for breast and mammary-specific gene sets. **(A)** GSEA MSigDB hits (based on number key in Table 1) for ATAC peaks in Hallmarks gene sets (left) and hits (based on letter key in Table 2) for breast and mammary-specific gene sets (right) in long-term TGFβ-induced and –withdrawn conditions. Gray = 0d; yellow = 10d +TGFβ; purple = 10d +TGFβ, 10d -TGFβ. **(B)** GSEA MSigDB hits for genes annotated to ATAC peaks with persistent chromatin alterations following TGFβ treatment and withdrawal in Hallmarks gene sets (top) and breast and mammary-specific gene sets (bottom)

**Supplementary Figure 4.**
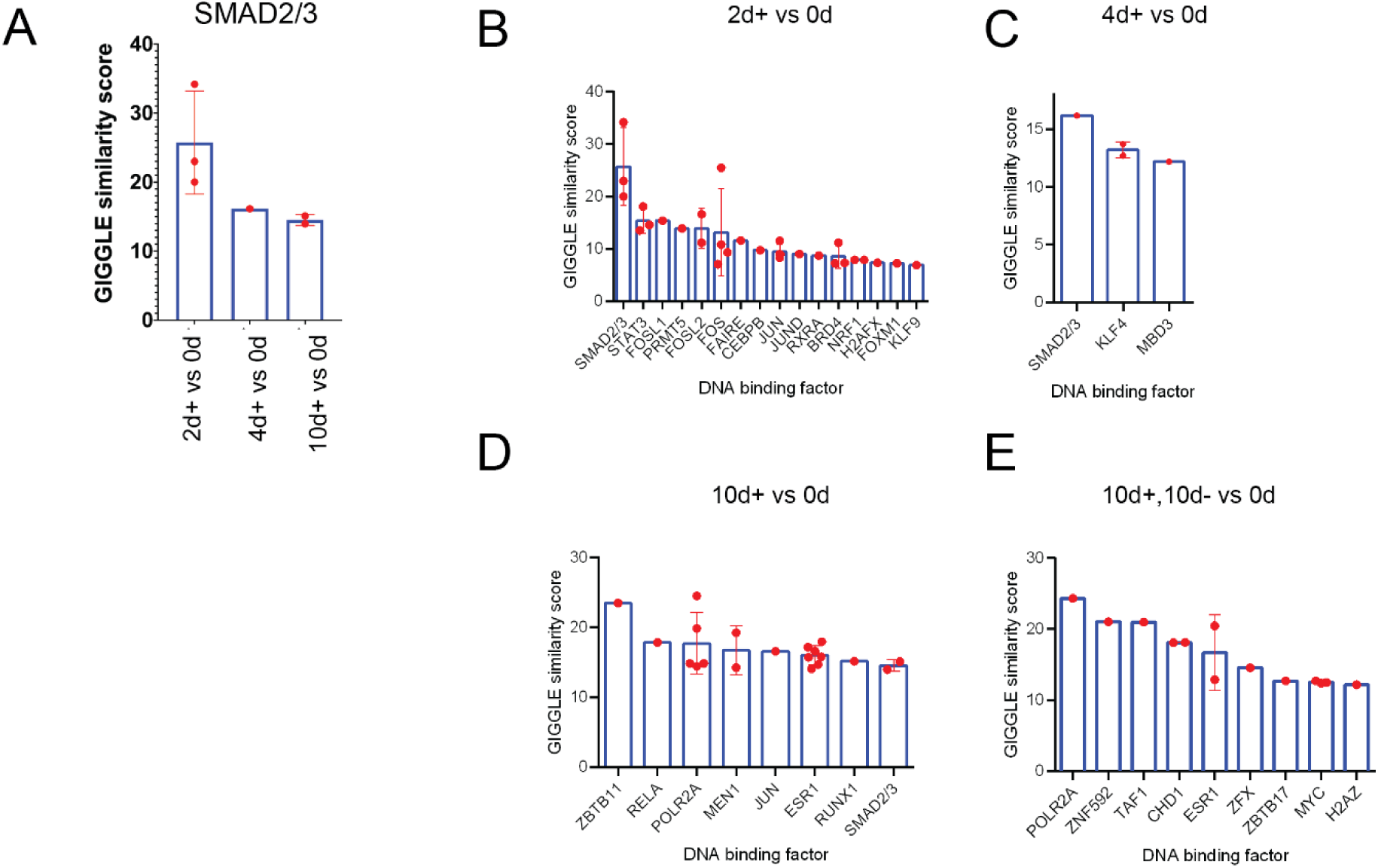
GIGGLE similarity scores of probable chromatin accessibility regulators at various stages of the EMT-MET spectrum in breast-specific cell lines. Peaks enriched at the indicated timepoints vs untreated cells were compared to genome interval files. **(A)** SMAD2/3-associated genome interval files show a diminished score as EMT progresses. **(B-E)** All scoring genome interval files for the indicated timepoint comparisons are shown. Data points refer to independent genome interval files queried by GIGGLE.

**Supplementary Figure 5.**
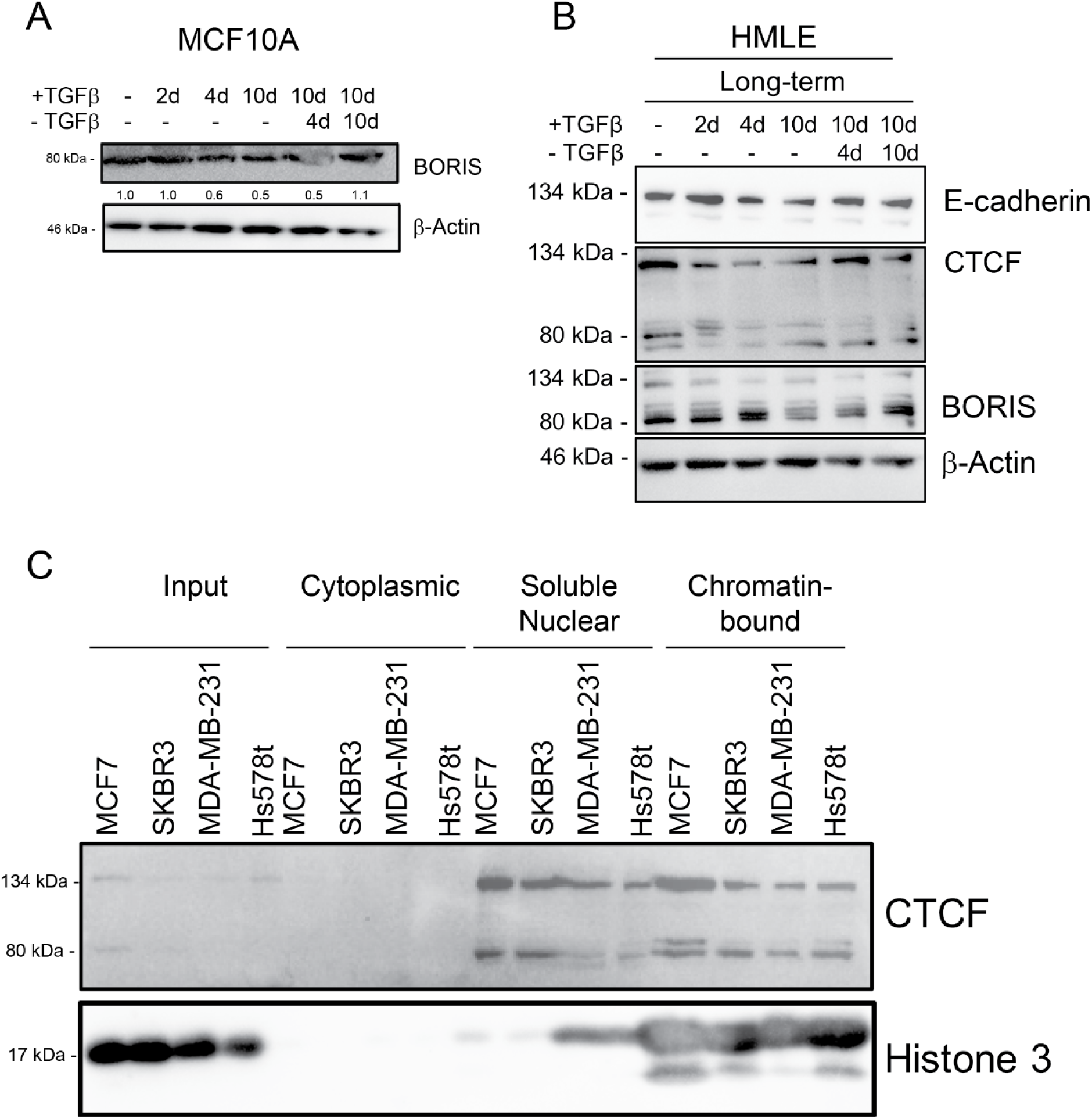
EMT suppresses total CTCF protein levels. **(A)** Western blot for BORIS in treated MCF10A cells. **(B)** Western blot for E-cadherin, CTCF, and BORIS in HMLE cell lines treated with (or withdrawn from) 5 ng/mL TGFβ at the indicated time. **(C)** Western blot for CTCF localization in specified cell fractions in the specified breast cancer cell lines.

**Supplementary Figure 6.**
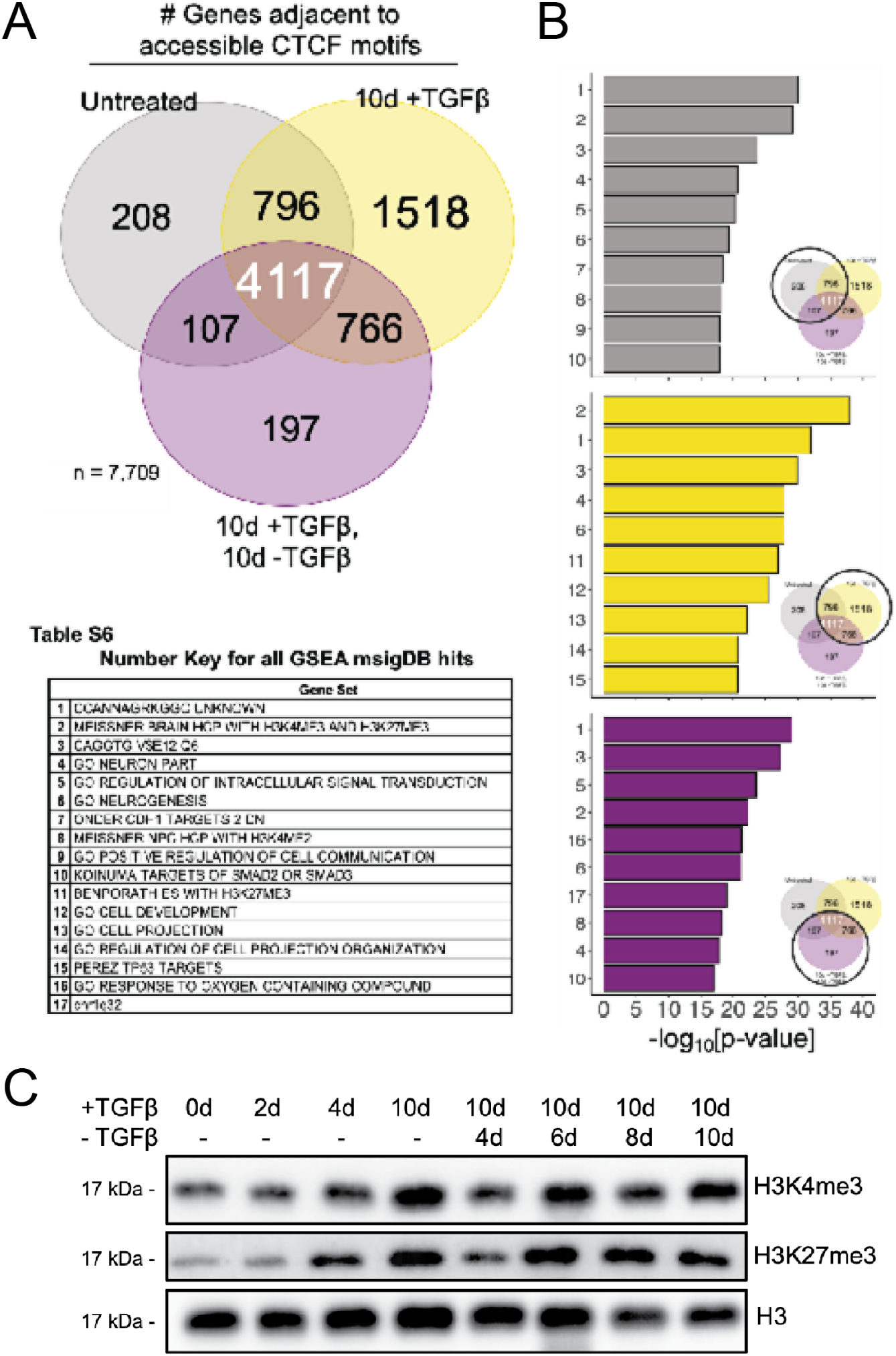
Genes near accessible CTCF motifs are enriched for bivalent, neuronal, and signal transduction genes. **(A)** Venn diagram representing the number of genes adjacent to accessible CTCF motifs at the indicated timepoints. **(Table S6)** GSEA MSigDB gene sets determined to be highly-enriched in accessible CTCF motifs. **(B)** GSEA MSigDB enrichment (based on number key in Table S6) for top-10 enriched gene sets by condition circled in venn diagram to the right. **(C)** Western blot for histone modification changes in treated MCF10A cells.

**Supplementary Figure 7.**
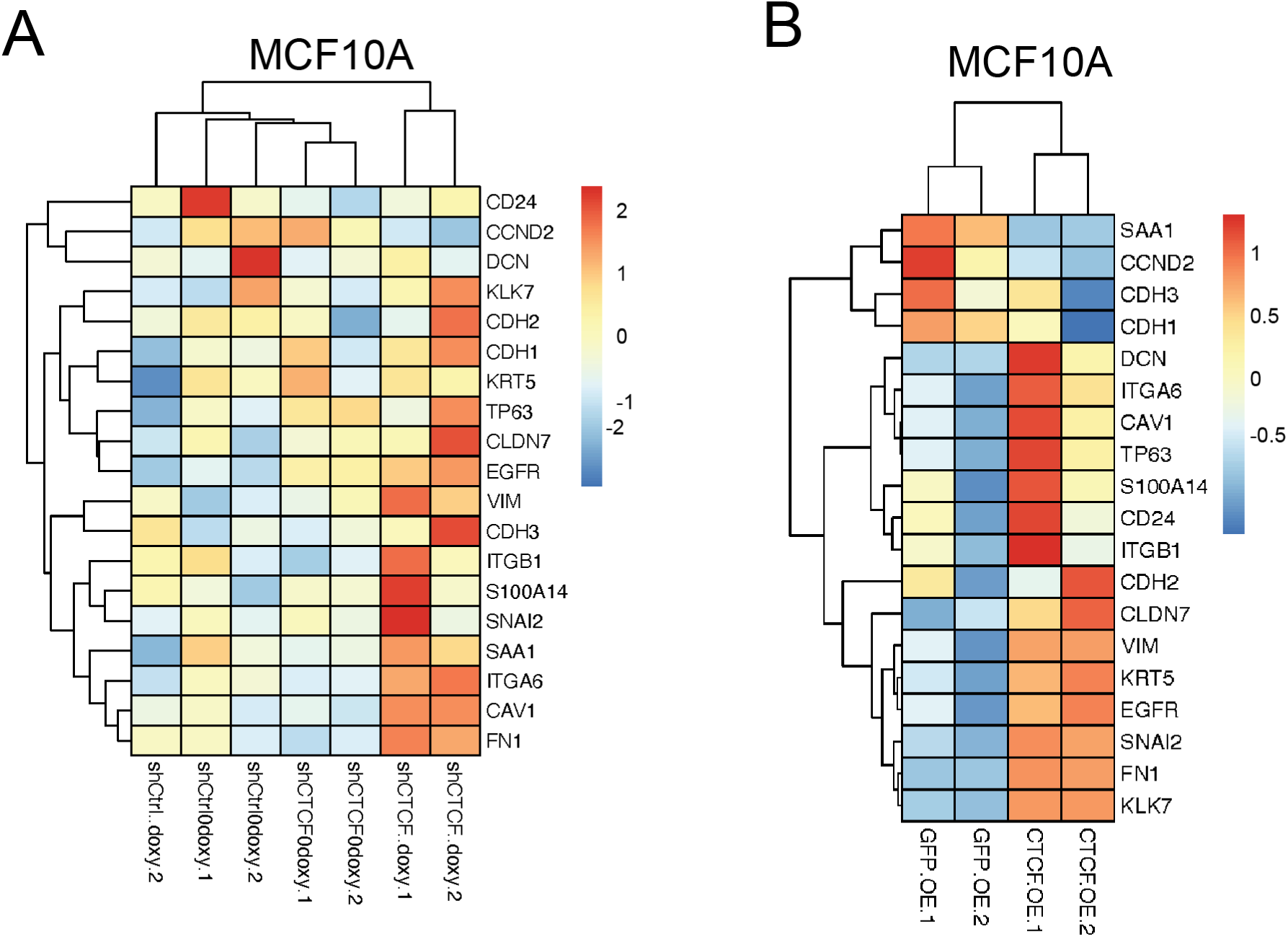
Change in gene expression upon CTCF manipulation in MCF10A cells. **(A)** Gene expression (normalized z-scores) was analyzed via Nanostring for indicated genes in shCtrl and shCTCF MCF10A cell lines with and without doxycycline. **(B)** Gene expression (normalized z-scores) was analyzed via Nanostring for indicated genes in GFP and CTCF overexpressing MCF10A cell lines.

